# Precision single-cell profiling of Circulating Tumour Cells: novel markers and data-driven characterization by CTCeek

**DOI:** 10.64898/2026.02.18.706522

**Authors:** Anna Terrazzan, Pietro Ancona, Francesca P. Carbone, Pachara Trevisan, Cristina Zuccato, Ewa A. Szymanek, Michał Szeląg, Federica Brugnoli, Anna Żaczek, Pawel Gaj, Michał Swierniak, Luana Calabrò, Chiara Agnoletto, Jeffrey Palatini, Nicoletta Bianchi, Renata Duchnowska, Elzbieta Senkus, Krystian Jazdzewski, Tomasz S. Kaminski, Stefano Volinia

## Abstract

Circulating tumour cells (CTCs) represent a minimally invasive method for monitoring cancer evolution in patients. CTCs are nowadays commonly isolated using antibodies against EPCAM protein. A key limitation regards the extent of EPCAM-negative CTCs, such as those that undergo EMT or whose tumour of origin is EPCAM-low or negative. We studied 3,302 RNA single-cell transcriptomes reported as CTCs in public repositories. Using copy number variation and cell type-specific markers, we discriminated *bona fide* CTCs from contaminating blood cells, often mislabelled as CTCs. The integration of *bona fide* CTCs and PBMCs, from multiple datasets, allowed us to identify novel markers, such as CLDN4, CLDN7, EFNA1 and TACSTD2 for epithelial CTCs, KCNK15 and LY6K for epithelial B CTCs, and ITGB4 for both epithelial B and mesenchymal CTCs. We revealed PODXL, AXL, CAV1, and TGM2 as markers of mesenchymal CTCs, which might be undetectable using anti-EPCAM antibodies, and TM4SF1 as universal marker, expressed in all CTC subclasses. Additionally, we found platelets to be physically associated with the epithelial A, but not with the epithelial B or the mesenchymal subtypes. Finally, we developed and implemented CTCeek, the first web-based and public reference tool that automatically annotates *bona fide* CTCs from scRNA-sequencing profiles.

## Introduction

Metastasis is the primary contributor to cancer mortality, accounting for an estimated 66%–90% of cancer-related deaths^(1)^. It is a multistep process that unravels through epithelial-to-mesenchymal transition (EMT), invasion, and colonization of distant organs^(2–5)^. Circulating tumour cells (CTCs), which are shed in the blood by the tumour, have the potential to form distant metastases^(6)^. A higher number of CTCs detected in cancer patients is associated with shorter survival^(7)^. Given their epithelial origin, CTCs’ fitness in circulation is thought to be diminished, due to phenomena such as anoikis, shear stress from blood flow, and the action of the immune system^(8)^. To endure the challenges in the circulation, CTCs may physically and functionally associate with platelets, erythrocytes, neutrophils, macrophages, natural killer (NK) cells, lymphocytes, endothelial cells, and cancer-associated fibroblasts^(9–10)^. Therefore, disclosing the deceptions that CTCs deploy in such an alien environment could lead to the design of novel therapeutic approaches^(11)^.

Cancer cells within a tumour can exist in different states, each associated with distinct functions, such as proliferation, differentiation, invasion, metastasis, and resistance to drugs^(12)^. Single-cell Next Generation Sequencing (scNGS) studies are profiling these diverse cell states^(13)^ to clarify the pathways which promote and sustain these functional states, regulating their transitions^(14,15)^, in relation with the tumour microenvironment^(16)^. We recently identified an epithelial CTC subpopulation with molecular characteristics akin to the trophectoderm, the embryonic layer that differentiates into invasive syncytiotrophoblast during human embryo implantation. Concurrently, using single-cell trajectory analysis, we showed that this CTC subpopulation originates from the lineage of metastatic tumours^(17)^.

CTC studies are still constrained by the availability of specific markers for their purification from blood, and current purification platforms limit clinicians and scientists from gathering complete information from the patients’ CTCs^(18,19)^. Despite the progress made in CTC research, each purification technique has its own limitations and biases, resulting in uneven purity of the enriched CTC population, which in most cases contains many non-CTC contaminants. Currently, there is no available tool, hitherto, which can reliably characterize and discern *bona fide* CTCs in single-cell RNA sequencing (scRNA-seq) datasets. Therefore, there is a pressing need to enhance comprehensive CTC harvesting from patients, thereby expanding the range of CTC purification platforms^(20)^. To address this, we collected and integrated over 3,300 scRNA-seq profiles of samples labelled as CTCs from publicly available databases^(21)^. These data have been analysed to develop a web-based tool which can help researchers in characterizing CTC populations from scRNA-seq data.

## Results

### Emergence of contaminating cells from the integration of candidate single-cell CTC profiles

We studied the transcriptomes of 3,302 putative CTCs obtained from 27 datasets from GEO or SRA public databases. Most of the RNA profiles were derived from blood of patients with breast cancer, but patients with pancreatic, prostate, lung, liver, gastric and colorectal cancers, or melanoma were also represented (Supplementary Table 1). Pre-processing and sample selection led to 1,230 quality control (QC)-passed single-cell RNA profiles for putative CTCs from the following cancer tissue types: breast (n= 1060), prostate (n= 105), gastric (n= 20), melanoma (n= 28), pancreas (n= 9), colorectal (n= 6), and lung (n= 2). The only two lung cancer-derived samples were excluded, because of low numerosity. The candidate CTCs are shown after integration and clustering in the Uniform Manifold Approximation and Projection (UMAP) of Fig. 1A. The summarizing score for epithelial carcinoma markers (keratins, like KRT8, KRT18 and KRT19) *vs* hematopoietic markers, i.e. protein tyrosine phosphatase, receptor type C (PTPRC/CD45), CD52, pro-platelet basic protein (PPBP) and platelet factor 4 (PF4), is displayed in Fig. 1B (red dots indicate cells of cancer origin, and blue dots those of hematopoietic lineage). The expression maps for the different markers, and for epithelial cell adhesion molecule (EPCAM) and vimentin (VIM), are shown in Supplementary Fig. 1A, B and C. Most cells on the UMAP plot had positive epithelial scores and were automatically annotated as breast cancer cells (Fig. 1C), therefore representing strong candidates for CTCs. Conversely, the blue dots in Fig. 1B indicated cells with negative epithelial scores and were also independently annotated as hematopoietic (e.g., macrophages, monocytes) or endothelial cells, namely non-CTC contaminants.

**Fig. 1.**
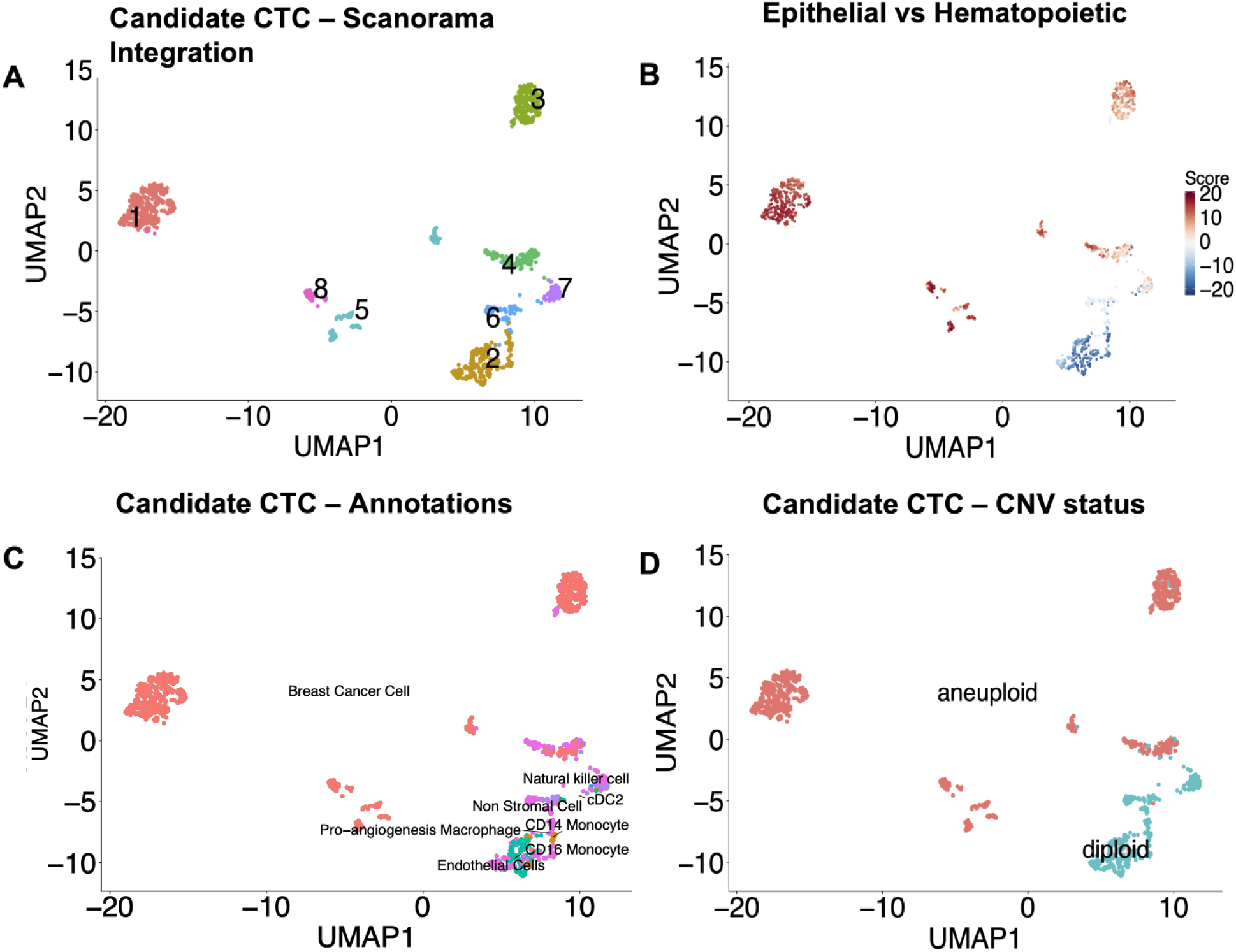
The Uniform Manifold Approximation and Projection (UMAP) representation of integrated putative CTCs. **(A)** Different colours indicate the clusters (1–8) based on marker expression and obtained after Scanorama integration and Leiden algorithm (with a resolution value K of 0.5). The genes involved in the cell cycle were removed prior to Scanorama integration to avoid clustering based on cell cycle. **(B)** The score derived from the expression of epithelial (red, positive) vs hematopoietic (blue, negative) markers. **(C)** Cell type automated annotation based on the expression profiles. **(D)** Copy number variations (CNVs) inferred for each putative CTC. CopyKat was used to identify cells with CNVs, indicated in red (aneuploid), while the azure dots are diploid cells.

The large KRT18-positive (Supplementary Fig. 1A) and VIM-positive (Supplementary Fig. 1C) cluster #3 stemmed from cancer cells and was thus constituted of CTCs with a mesenchymal state. Notably, while EPCAM (Supplementary Fig. 1B) was clearly restricted to cancer-derived KRT8/KRT18-positive CTCs, VIM expression was shared by both CTCs and hematopoietic cells.

From this analysis it emerged that a sizeable portion of the putative CTCs was composed of hematopoietic, or more accurately, blood-borne, non-CTC contaminants. To orthogonally validate such separation of strong candidate CTCs from contaminating non-cancerous blood-borne cells (either PTPRC/CD45 positive or negative), we then inferred copy number variations (CNVs). Cancer cells often bear multiple and extensive CNVs in their chromosomal repertoire resulting in aneuploidy, which is essentially absent in non-cancer cells^(19)^. Reassuringly, in the UMAP plot (Fig.1D) there was an almost complete overlap between aneuploid (highly likely to be cancer) cells, and cells with epithelial features (Fig. 1B). We thus assumed that the KRT-positive, CD45-negative (Supplementary Fig. 1D), PPBP-negative (Supplementary Fig. 1E) and aneuploid cells (Fig. 1D, red dots) were CTCs; and only clusters containing such cells were deemed to contain *bona fide* CTCs. Finally, the cell cycle stages were inferred for each cell and shown in Supplementary Fig. 2; strikingly, only CTCs, but not the contaminant cells, were actively engaged in the cell cycle.

### Pure CTCs identification by integrative analysis of putative CTCs and PBMCs

Our primary goal was to identify novel and informative markers to aid in the detection, purification, and targeting of CTCs within the bloodstream. Thus, we needed to pinpoint with high confidence the *bona fide*, or “pure”, CTCs, based on the results of the integrated CTC datasets shown above. Our analysis had shown that only a fraction of the cells labelled as CTCs in the public databases were likely of cancer origin, while the remaining portion was of hematopoietic or endothelial origin. To definitely establish which candidate CTCs were from blood rather than cancer sources, we integrated the CTCs (n= 1228) with scRNA-seq profiles of PBMCs (n= 10,434)^(22)^. Since there were major differences in cell cycle activation throughout the various cell types, as shown in the earlier CTC-focused analysis (Supplementary Fig. 2), prior to integration, we removed the cell cycle genes from the scRNA-seq profiles. While this choice might hinder some biological meaning from the integrated profiles, it clustered together cells based on their types, regardless of mitotic activation.

The first step in our quest was tackling the cellular complexity by overlapping the Scanorama cluster map (Fig. 2A) with the CTC annotation from the CTC-only integration performed earlier (Fig. 2B and C), and with the PBMC annotation (Fig. 2D). Most CTC contaminants were mapped to clusters #25 and #34. As expected, the contaminant cells did not have a detectable expression of CD45/PTPRC (Fig. 2E), and for this reason, were wrongly selected together with *bona fide* CTCs, but they were also mainly diploid, a characteristic akin to non-cancer cells (Fig. 2C). Differential expression analysis of the Scanorama clusters identified endothelial and erythroid genes, together with platelet restricted genes (PBPP, PF4), as markers (https://panglaodb.se/markers.html) for the contaminant cells (Fig. 2F and Supplementary Table 2). The boundary between the candidate CTC cluster #32 and the putative contaminant cluster #34 was fuzzy, because of a decreasing platelet content in the CTCs (not shown). Interestingly, besides two platelets, seven nucleated cells from the PBMC dataset were mapped to the CTC contaminants. This finding confirmed that there are rare, nucleated blood-borne cells (7/10434= 0.0006) that can likely be captured in a liquid biopsy along with CTCs, thus resulting in an erroneous classification.

**Fig. 2.**
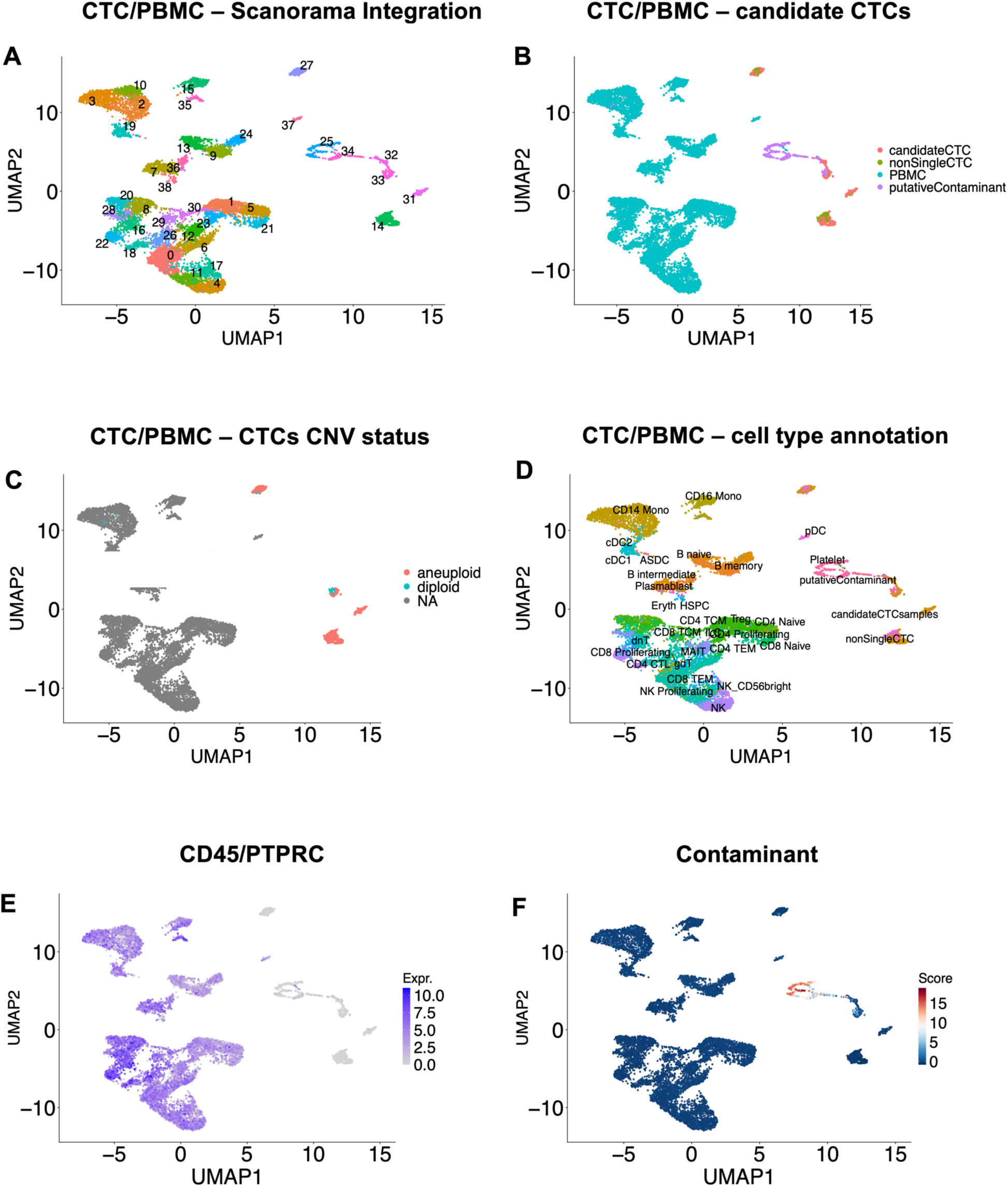
Integration of single-cell RNA-seq CTC and PBMC datasets (Step I). **(A)** Scanorama integration of QC-passed CTC and PBMC single-cell datasets, indicated by different colours corresponding to Leiden clusters (K= 2); **(B)** Scanorama UMAP shows PBMCs (azure), used as reference for blood cells, bona fide CTCs (single cells in red, and CTC clusters in green), with non-cancer diploid contaminant cells (violet) that co-purified along with CTCs; **(C)** CNV status for putative CTCs; **(D)** Cell type annotations for PBMCs. **(E)** Expression of CD45/PTPRC, a PBMC marker; **(F)** The contaminant score (obtained as the mean expression of the markers for endothelial cells and platelets).

We then evaluated the presence of contaminant cells across each original CTC dataset. Nearly all CTC datasets contained non-CTC cells, but the rate of these false positives changed vastly among datasets, and technical platforms, ranging from 1% up to 100%, with a median contamination rate of 54% (Supplementary Table 3). Microfluidics- and/or label-free CTC enrichment typically resulted in low recovery of *bona fide* CTC and high co-purification of non-CTC cells. Only the systems employing positive selection with anti-EPCAM, or other CTC-specific antibodies, achieved false positive rates below 10%.

We excluded the samples from the two contaminant clusters (#25 and #34) and finally obtained a large *bona fide* and pure CTC population (n= 900). We then performed the integration of the bona fide CTCs with PBMCs to obtain refined embedding and clusters (Fig. 3A). The keratins-positive CTCs in this integration were divided into three major groups and corresponded well with the cancer cells (Fig. 3B and C). All CTC clusters expressed high levels of EPCAM (Fig. 3D), except for the VIM-positive cluster #27, deemed thereafter as mesenchymal (Fig. 3E). The heterogeneous CTC group of three contiguous clusters (#28, #38, and #47) was interestingly bordering with the hematopoietic stem and progenitor cell (HSPC) (cluster #48). Cluster #38 contained the prostate CTCs (Fig. 3B) and its tumour of origin well correlated with the expression of kallikreins related peptides, KLK2/KLK3 (Fig. 3F), and androgen receptor (AR) (Supplementary Fig. 3). As expected, the mesenchymal CTCs, expressed high levels of genes involved in EMT, such as ZEB1/ZEB2 and Snail Family Transcriptional Repressor (SNAI) 1 and 2 (Supplementary Fig. 4). Over-expressed genes for membrane-bound proteins in the epithelial cluster #8 included Lymphocyte Antigen 6 Family Member K (LY6K), as expected for the epithelial B CTCs^(17)^, in contrast with the remaining contiguous epithelial A CTC clusters (#28, #38, and #47).

**Fig. 3.**
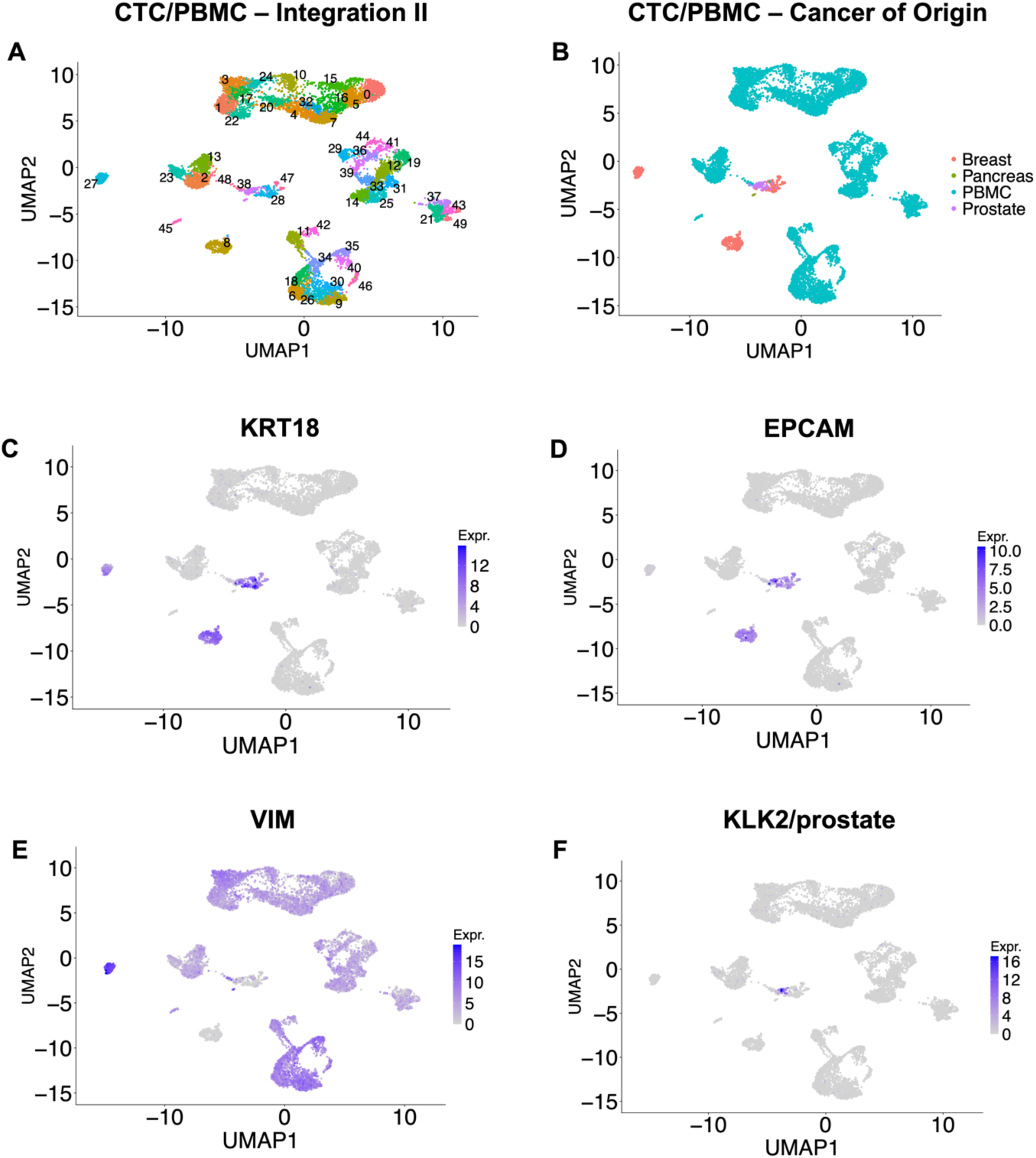
Integration of single-cell RNA-seq CTC and PBMC datasets after the removal of CTC contaminants (Step II). **(A)** Scanorama integration of QC-passed CTC and PBMC single-cell datasets, indicated with 28 different colors corresponding to Leiden clusters (K= 2); **(B)** Scanorama UMAP shows the cancer of origin for the CTCs (breast, prostate, or pancreas) in the PBMC background (azure); **(C)** Expression of KRT18, a cancer marker; **(D)** EPCAM, an epithelial marker. **(E)** VIM, a mesenchymal marker; both PBMCs and CTC cluster #27 are positive; **(F)** KLK2, a prostate marker.

### Identification of novel markers for the enrichment of CTCs from blood

The identification of *bona fide* CTCs finally allowed us to search for novel CTC markers as alternatives to the current EPCAM gold standard. At first, we looked for pan-CTC markers, i.e. markers capable of selecting CTCs regardless of their status (epithelial/mesenchymal), or tissue of origin. To obtain robust markers, we compared CTCs *vs* PBMCs using both single-cell and pseudo-bulk RNA profiles. The newly discovered markers are shown in Table 1, labelled with an asterisk, together with other genes expressed in CTCs but not in PBMCs. Notably, the genes for membrane-associated proteins such as Tumour-associated Calcium Signal Transducer 2 (TACSTD2), Syndecan-4 (SDC4), Claudin 7 (CLDN7) and Transmembrane 4 L six family member 1 (TM4SF1) had AUC levels as high as those of known epithelial markers KRT7, KRT8, KRT18, KRT19, and EPCAM.

**Table 1.** The marker genes over-expressed in CTCs when compared with PBMCs.

| Gene | AUC<br>(single-cell) | Avg log2<br>FC<br>(single-cell) | Adjusted<br>p-value<br>(single-cell) | AUC<br>(pseudo-bulk) | Avg log2<br>FC<br>(pseudo-bulk) | BH<br>adjusted<br>p-value<br>(pseudo-bulk) |
| --- | --- | --- | --- | --- | --- | --- |
| KRT18 | 0.99 | 18.4 | 1.00E-03 | 1.00 | 7.6 | 0.0004 |
| TOMM6* | 0.99 | 14.0 | 1.00E-03 | 1.00 | 6.6 | 5.65E-07 |
| KRT8 | 0.98 | 17.3 | 1.00E-03 | 1.00 | 7.1 | 0.0004 |
| CTTN* | 0.98 | 11.1 | 1.00E-03 | 1.00 | 6.5 | 0.0002 |
| MLPH* | 0.97 | 23.8 | 1.00E-03 | 1.00 | 7.5 | 0.0002 |
| KRT19 | 0.96 | 21.6 | 1.00E-03 | 1.00 | 8.2 | 1.36E-06 |
| TACSTD2 *<br>(TROP2) | 0.96 | 31.5 | 1.00E-03 | 1.00 | 7.3 | 1.52E-06 |
| CLDN7* | 0.96 | 10.1 | 1.00E-03 | 0.99 | 6.1 | 0.0004 |
| SDC4* | 0.95 | 12.2 | 1.00E-03 | 1.00 | 6.7 | 0.0004 |
| EPCAM | 0.95 | 12.2 | 1.00E-03 | 0.99 | 6.4 | 1.60E-05 |
| TM4SF1* | 0.94 | 19.8 | 1.00E-03 | 1.00 | 8.1 | 5.65E-07 |
| KRT7 | 0.94 | 15.1 | 1.00E-03 | 1.00 | 7.2 | 2.13E-05 |
| SDC1* | 0.94 | 13.2 | 1.00E-03 | 0.99 | 6.8 | 1.56E-06 |
| NME1-NME2 | 0.92 | 17.4 | 1.00E-03 | 1.00 | 6.9 | 5.65E-07 |
| CLDN4* | 0.91 | 17.9 | 1.00E-03 | 0.99 | 7.2 | 5.89E-05 |

Single cell AUC>0.9. Benjamini Hochberg (BH) bulk adjusted p-value <0.05; bulk average log2 Fold Change (avg log2 FC) >6. Novel CTC markers are labelled with asterisk (\*).

The differentially expressed genes for the two epithelial CTC subgroups, alone or in combination, are listed in Supplementary Tables 4-6.

### Epithelial versus mesenchymal CTCs: emergence of specific markers

We then evaluated the expression of the newly identified markers also on the previously excluded, non-pure, candidate CTCs (Fig. 4). This comprehensive analysis allowed us to measure the new CTC markers on a wider, albeit somewhat contaminated, CTC population. Looking across the diverse cancer types, most CTC markers were restricted to breast and/or prostate, but not all. For example, TM4SF1, a membrane transporter protein and cancer antigen, was detected in CTCs from five out of six malignancies, with melanoma being the only exception (Fig. 4A and B). In contrast, the gold standard for CTC detection, EPCAM, was expressed only in three cancer types: breast, colorectal, and prostate, but was absent in pancreatic and stomach cancers (Fig. 4C and D). Conversely, the other epithelial CTC markers, TACSTD2 and CLDN7, were more expressed in breast and prostatic cancer derived cells (Fig. 4E-H). We then compared the newly identified CTC markers with current gold-standard EPCAM, across each CTC cluster and PBMC cell type (Supplementary Fig. 5A-D): ephrin A (EFNA1), a secreted protein anchored to the membrane^(23)^, was a marker for epithelial CTCs, but not mesenchymal CTCs (Supplementary Fig. 5B); TM4SF1 and CLDN7 were confirmed to be CTC specific (including the mesenchymal subtype) and negative in all PBMC cell types (Supplementary Fig. 5C and 5D). The epithelial B subclass had a very distinctive RNA profile with several specific genes, such as SP6 and LHX1 transcription factors, WNT3A and BMP7 growth factors, and KCNK15 and LY6K membrane proteins (Supplementary Fig. 5E and F). The expression profiles of single-cell mesenchymal CTCs were also very distinctive.

**Fig. 4.**
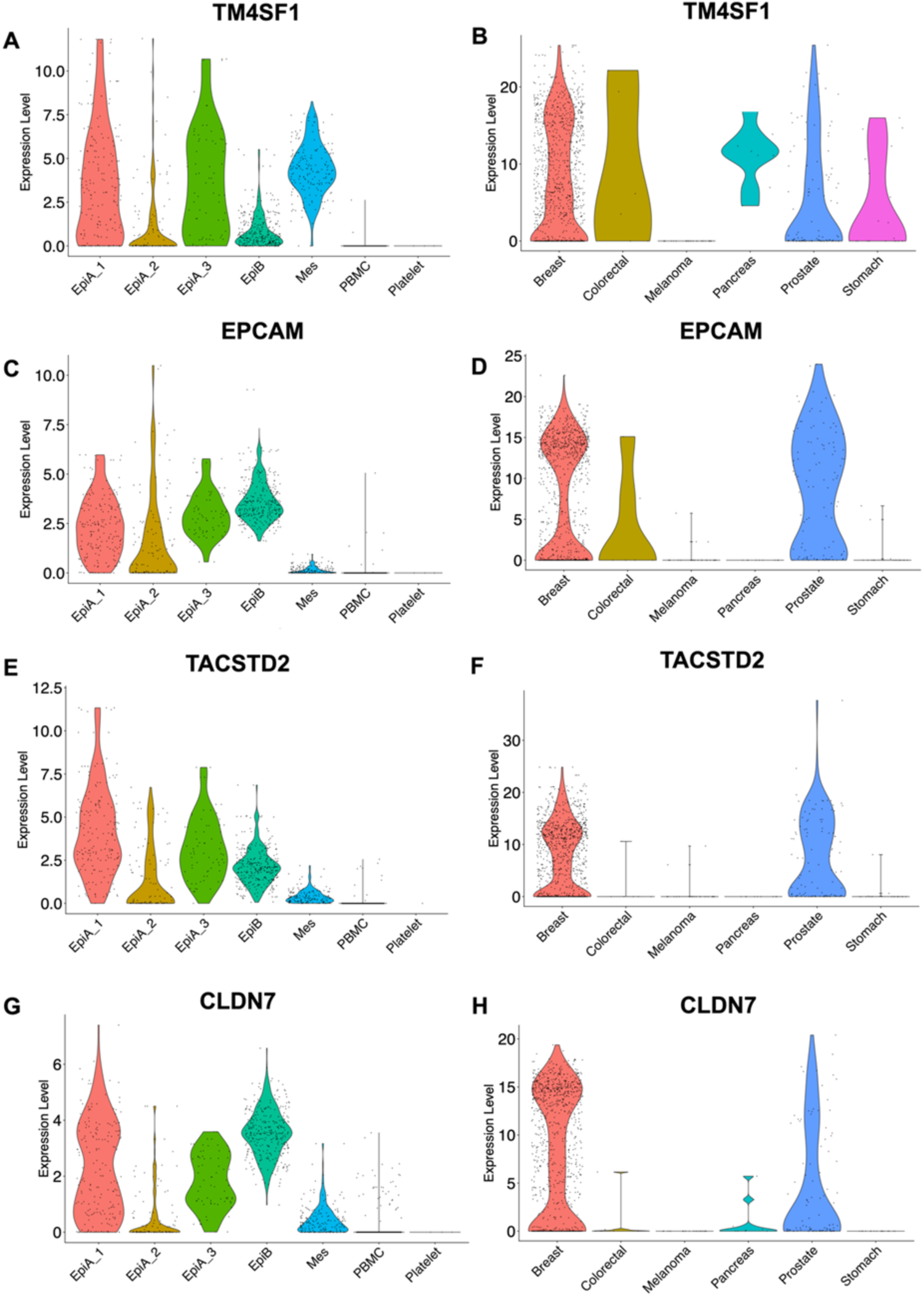
Violin plots of the novel markers for CTCs. The left column shows the expression across CTCs’ subgroups and contaminant cells, whereas the right column shows the expression across different cancer origins.

Unexpectedly, we identified several mesenchymal CTC-restricted mRNAs (Table 2), which could be used for both detection and purification, as some of their products are membrane proteins with extracellular domains. These novel markers included epithelial membrane protein 1 (EMP1), transglutaminase 2 (TGM2), AXL, caveolin-1 (CAV1) and podocalyxin (PODXL).

**Table 2.** The genes over-expressed in mesenchymal CTCs.

| Gene | AUC<br>(single-cell) | Avg log2FC (single-cell) | Adjusted<br>p-value<br>(single-cell) | AUC<br>(bulk) | Avg log2<br>FC (bulk) | BH<br>Adjusted<br>p-value (bulk) |
| --- | --- | --- | --- | --- | --- | --- |
| CAV1 | 1.00 | 9.48 | 1.00E-03 | 1.00 | 8.73 | 4.44E-05 |
| COL6A1 | 1.00 | 15.38 | 1.00E-03 | 1.00 | 9.16 | 5.97E-04 |
| EMP1 | 1.00 | 9.85 | 1.00E-03 | 1.00 | 6.90 | 1.51E-03 |
| AXL | 1.00 | 4.03 | 1.00E-03 | 1.00 | 6.29 | 1.71E-03 |
| TGM2 | 1.00 | 17.05 | 1.00E-03 | 1.00 | 8.60 | 2.26E-03 |
| RAI14 | 0.99 | 6.47 | 1.00E-03 | 1.00 | 6.16 | 4.83E-04 |
| COL13A1 | 0.99 | 8.81 | 1.00E-03 | 1.00 | 7.83 | 5.44E-05 |
| MYEOV | 0.99 | 10.56 | 1.00E-03 | 1.00 | 7.30 | 4.44E-05 |
| LAMA5 | 0.99 | 11.28 | 1.00E-03 | 1.00 | 6.07 | 2.50E-03 |
| IGFBP4 | 0.99 | 16.24 | 1.00E-03 | 1.00 | 6.89 | 2.20E-03 |
| EHD2 | 0.99 | 9.00 | 1.00E-03 | 1.00 | 6.67 | 1.61E-03 |
| FN1 | 0.98 | 11.23 | 1.00E-03 | 1.00 | 8.52 | 3.14E-03 |
| PODXL | 0.98 | 10.04 | 1.00E-03 | 1.00 | 7.40 | 4.83E-04 |
| DLC1 | 0.98 | 7.91 | 1.00E-03 | 1.00 | 7.30 | 4.14E-04 |
| IGFBP6 | 0.98 | 8.69 | 1.00E-03 | 1.00 | 8.03 | 8.36E-05 |

bulk adjusted p-value <0.05; bulk avg log2 FC >6

The violin plots for CAV1, AXL, and TGM2 are shown in Fig. 5, alongside VIM, the current non-specific marker for mesenchymal CTCs, which is also highly expressed in both PBMCs and platelets.

**Fig. 5.**
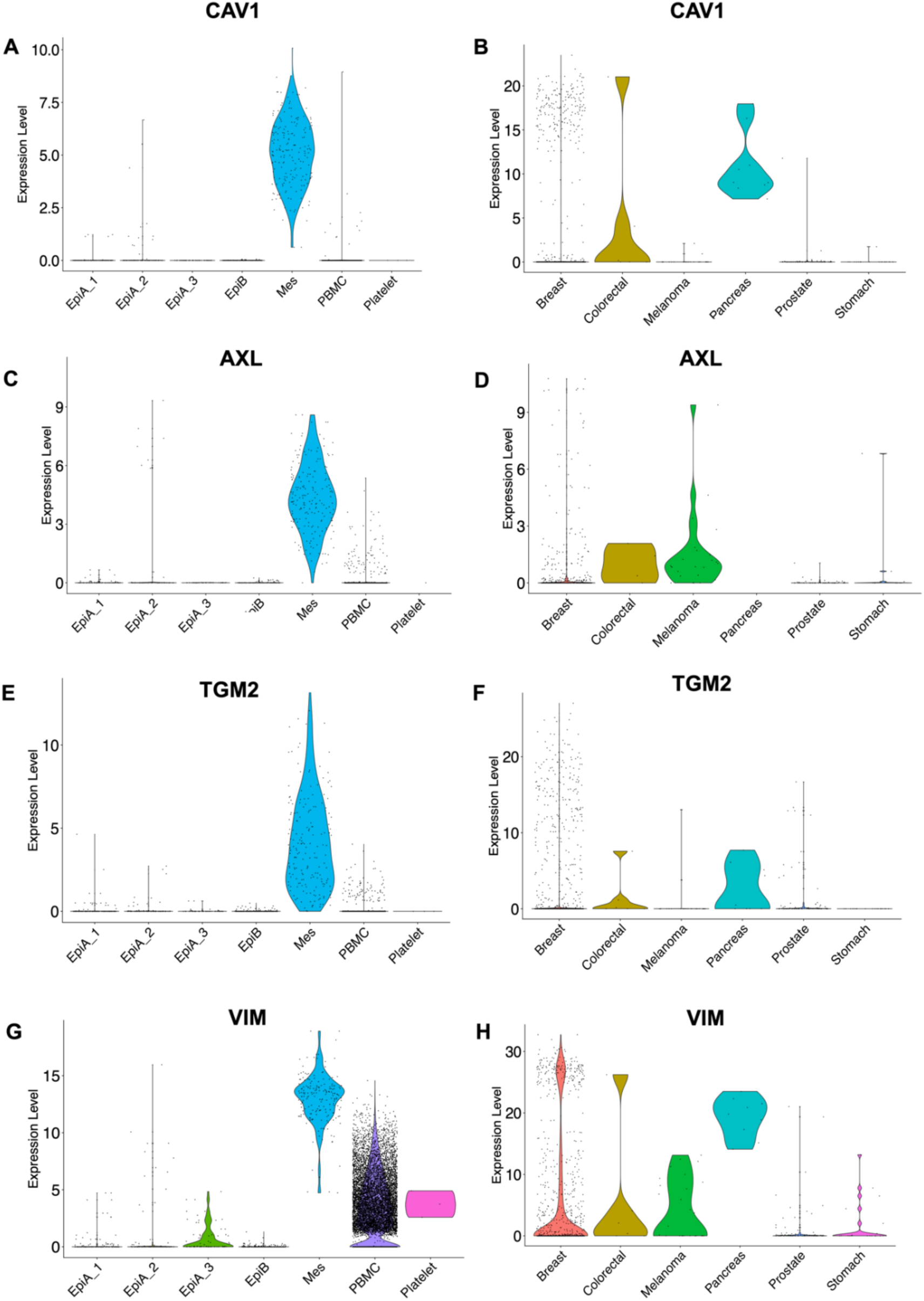
Violin plots of novel markers for mesenchymal CTCs. Left plots show the expression across cell types, while the right column shows the expression across the different cancer origins.

The detailed expressions of the mesenchymal markers described above are shown in Supplementary Fig. 6A-E. It was apparent from our initial work that cell cycle engagement was varied in extent within the CTC subclasses. The only two clusters with mitotically active CTCs were those for the epithelial B and the mesenchymal CTCs; the integrin subunit beta 4 (ITGB4), which plays a critical role in the hemidesmosome of epithelial cells, and is likely involved in the biology of invasive carcinoma^(24)^, was the highest gene expressed in those CTC subclasses (Supplementary Table 7 and Supplementary Fig. 6F).

Figure 6 shows the most prominent CTC markers we identified in this study, alongside KRT18 and EPCAM as references. Interestingly, platelets were abundant in the epithelial A CTCs but absent in the two mitotically active CTC subpopulations: epithelial B and mesenchymal CTCs.

**Fig. 6.**
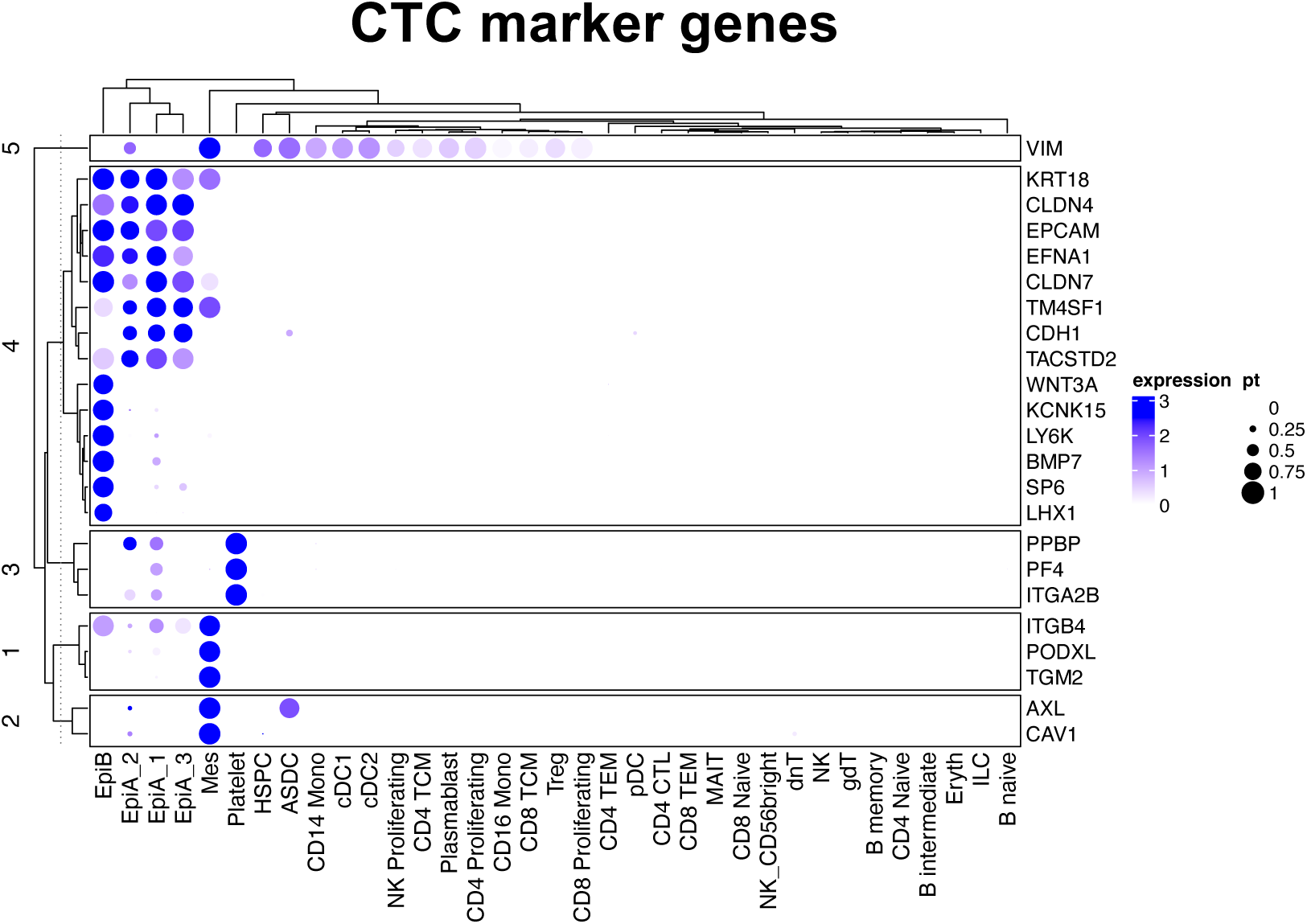
The novel markers for CTCs compared with PBMC cell types. Clustered dot-plot shows the genes over-expressed in diverse CTC subgroups. CTC clusters are indicated together with the hematopoietic cell types in the PBMCs. The radius of the circle is proportional to the percentage (pt) of positive cells in each lane.

### Negative selection of PBMCs and depletion of contaminant blood cells after CTC enrichment

A common step in the purification of CTCs from patients’ blood is the negative selection of hematopoietic cells. Usually, an anti-CD45 (PTPRC) antibody is applied to remove the excess of PBMCs from the CTC sample^(6)^. Indeed, when PTPRC/CD45 was tested for differential expression in PBMCs compared to CTCs (Supplementary Table 8), its ROC AUC was very high (0.91). Nonetheless, another marker, CD52 had an AUC even higher than that of PTPRC/CD45 (0.95), making it another suitable target for the negative selection of PBMCs. Also, CD48 and CD37 shared very high AUCs, as confirmed in the pseudo-bulk analysis, and the characteristics to serve as targets for physical removal of PBMC. It was apparent that PTPRC alone cannot cover all different PBMC cell types, and a combination of antigens, including CD48, CD52, CD37, and CXCR4 might be required for the depletion of most PBMCs (Supplementary Fig. 7).

Perhaps surprisingly, a large portion of blood-derived CTC contaminants was PTPRC-negative, with endothelial cells and platelets comprising the majority of false CTCs. Such residual contaminants could be distinguished using markers reported in Supplementary Table 2, such as: Transmembrane Protein 40 (TMEM40), Selectin P (SELP), C-Type Lectin Domain Family Member B (CLEC1B), Platelet Endothelial Aggregation Receptor-1 (PEAR1), Glycoprotein IX Platelet (GP9), and the integrin subunit alpha 2b (ITGA2B).

Finally, we also addressed the issue facing the usage of AXL as a positive marker for the selection of mesenchymal CTCs. AXL is also expressed in a rare population of dendritic cells, AXL^+^/SIGLEC6^+^ dendritic cells (ASDC) (Fig. 6). Thus, we leveraged our CTC/PBMC integrated dataset to identify possible markers for the negative selection of ASDCs. Genes for hematopoietic-restricted membrane proteins TYROBP (monocyte lineage) and HLA-DRB1 (antigen-presenting cells) were highly expressed in ASDCs but not in CTCs (Supplementary Table 9). Therefore, these last two proteins can also be used for the negative selection of ASDCs, as well as PBMCs in general.

### CTCeek: a web-based public tool to identify CTCs in scRNA-sequencing profiles

We have shown here that all CTCs datasets we studied, generated using different CTCs isolation methods^(20)^, contained contaminant blood-borne cells. To help with the identification of *bona fide* CTCs we therefore developed CTCeek, a web-based app that automatically identifies the CTCs present in scRNA-seq profiles. CTCeek maps the query cells onto the PBMC/CTC reference’s UMAP embedding space, allowing for their direct visualization on the reference atlas. To test the functionality and performance of CTCeek, we used both negative (PBMC) and positive (CTC) controls. The formers consisted of PBMCs from healthy individuals: the pbmc3k (n= 2,700 cells), and the Broad Institute’s 44k PBMC datasets^(22)^ (n= 44,433 cells) from which cells used in the reference were removed (n= 9922 cells). As positive CTC controls, we used samples (GEO entries: GSE255889 and GSE295441)^(25,26)^ that, again, were not in the reference dataset. The CTC validation dataset contained contributions from patients with various types of metastatic cancers, resulting in 136 putative CTCs (Supplementary Table 10), with 88 of them inferred as aneuploid by both CopyKAT and SCEVAN^(27,28)^. It must be noted that while aneuploid cells are very likely CTCs, we also expected a small but sizeable portion of diploid CTC as well (i.e. true CTCs for which the aneuploidy test was negative). CTCeek predicted 93 *bona fide* CTCs (out of 136) and 82 *bona fide* CTCs out of the 88 aneuploid cells (details in Supplementary Table 11). CTCeek called the six aneuploid contaminant cells as hematopoietic stem and progenitor cells (HSPCs) (n= 4) or as platelets (n= 2). Although SCEVAN and CopyKAT had predicted these six cells to be aneuploid/tumour, gene marker analysis confirmed the CTCeek classification. Specifically, the four “aneuploid” HSPCs expressed CD34 and FLT3, while the “aneuploid” platelets expressed PPBP, as shown in Supplementary Fig. 8. Regarding the benchmark with the “pbmc3k” dataset, the tool showed 99.96% of specificity and a corresponding false positive rate (Type I error) of 0.037% (see Methods). Concerning the Broad Institute’s PBMCs dataset, including scRNA-seq profiles obtained with several UMI-based technologies (CEL-Seq2, 10x Chromium, Drop-seq, Seq-Well, and inDrops), CTCeek likewise performed optimally, obtaining a specificity of 99.99% and a false positive rate of 0.01%. Indeed, our tool identified only 1 CTC (false positive) out of a total of 34,511 PBMCs. The statistics summary for this benchmark is reported in Supplementary Table 11.

## Discussion

CTCs and circulating tumour DNA (ctDNA) are emerging as two main actors in the minimally invasive study of cancer evolution. Both players have been the subject of intensive technical improvements, and next-generation sequencing has been shown to deliver breakthroughs in the molecular characterization of ctDNA, as well as of CTC RNA repertoire. Our goal in this study was to identify novel molecular markers to improve the isolation of CTCs, particularly for the currently under-studied EPCAM-low CTCs. We leveraged our work on the several single-cell level CTC studies published in recent years. Because CTCs are known to be present in two “flavours”, epithelial and mesenchymal, we attempted to identify both pan-CTC or status-specific markers, particularly for the elusive EPCAM-low mesenchymal state. Meanwhile, our study also revealed some interesting findings. The first one was that essentially all aneuploid cells were *bona fide* CTCs and vice versa. As expected, aneuploid cells were epithelial (cancer-derived cells) and expressing cytokeratins. Consequently, diploid cells in the CTC fractions, that were formerly labeled as cancerous in their original studies, were indeed mostly false positives. This clear separation could emerge only when an integrative multi-study analysis was performed, as often the narrowness, or low statistical power, of each study does not allow a robust statistical analysis of inferred CNV status. A second finding was that a large fraction of CTCs was engaged in the cell cycle, in contrast to the PBMCs and the contaminant cells. Finally, CD45-negative, and diploid blood cells, commonly enriched by size selection, were heavily contaminating the CTC population.

We described a collection of novel markers that expands the horizon of CTC beyond the one defined by the EPCAM paradigm. In fact, we identified both pan-CTC markers, and markers restricted to either epithelial or mesenchymal state. Three of the novel markers we identified, TACSTD2, CLDN4 and AXL have also been implicated in CTCs by other investigators. Liao *et al*.^(29)^ recently described TACSTD2 (TROP2) as a marker for “epithelial mesenchymal CTCs”, in triple-negative breast cancer; in our much larger CTC population, TACSTD2 was highly expressed in CTCs, less in those with mesenchymal status, and was present in breast and prostate derived cells. Strikingly, ADC therapy using antibodies against TACSTD2 (TROP2) is already used to treat metastatic TNBC^(30)^, in patients that are PD-L1 low. Targeting cells expressing TACSTD2 led to significantly longer progression-free survival than chemotherapy among patients with advanced triple-negative breast cancer who were not candidates for treatment with PD-1 or PD-L1 inhibitors. Finally, Chai *et al*. showed that the number of CTCs was positively correlated with the CLDN4’s expression in the primary breast tumours^(31)^. Bardol *et al*. identified AXL-positive CTCs in 7 out of 60 patients using the CellSearch*^®^*system. They propose that those AXL-positive and EPCAM-selected CTCs have just been ongoing EMT^(32)^, in agreement with our finding of AXL in the mesenchymal CTCs. We also shed additional light on the role of platelets: these small cell fragments are important companions of CTCs and are thought to be involved in the invasive process. It has been proposed that platelets support tumour metastasis and have a crucial role in the progression of cancer^(33)^. Our study confirmed the association between thrombocytes and CTCs, but restricted to epithelial A, and not to epithelial B nor mesenchymal CTC subclasses. The two latter, mitotically active CTC subclasses, maybe be independent from platelet protection.

There is another hurdle in obtaining highly enriched CTCs using microfluidics or other physical purification platform. We have shown that “false” CTC contaminants were very frequent upon microfluidic enrichment, even in combination with anti-CD45 depletion. Most of the CTC datasets available form GEO contained some PTPRC-positive cells but, unexpectedly, a considerable fraction of blood-derived CTC contaminants lacked PTPRC expression. Within our integrated datasets, we could precisely compute the number of expected CD45 false positive CTCs, obtained by size selection of human blood. Standard label-free and microfluidics-based CTC purification from a 5 mL blood sample can result in up to 6,000 CD45-negative blood-borne non-CTC cells, at least in healthy blood. Consequently, effective depletion can be achieved using antibody-coated beads targeting our newly identified specific markers: SELP, TMEM40, CLEC1B, PEAR1, GP9, and ITGA2B. Antibodies targeting these molecules, in conjunction with the negative selection of PBMC, could then be employed to enrich CTCs without using CTC-targeted antibodies. Such a platform would ensure an unbiased representation of the whole CTC range in cancer patients, regardless of their subgroups, which might be relevant in under-studied cancer types, or individual patients. These will allow to overcome the limitations of current strategies (relying solely on established markers, i.e. EPCAM) which often produce fewer desirable results.

The limitations in our study were various and mostly related to the current technologies and platforms. Although scRNA-seq has now reached a degree of maturity, we had to discard many samples to enable high quality control and high molecular complexity levels. Additionally, we obtained the CTC data from several different projects and had to rely on integration to study the datasets in a monolithic fashion. Furthermore, most, albeit not all, CTCs included in this study were derived from patients with breast cancer, therefore our results might not robustly apply to carcinomas other than breast cancer. Again, many of the single-cell profiles were from EPCAM-selected CTCs and, although we also identified a sizable population of EPCAM-low mesenchymal CTCs, this bias might have led to a skewed representation of the CTC spectrum. Finally, RNA and protein levels are not always correlated, and the abundance of markers should be verified by conventional immunofluorescence or with single-cell proteome analysis methods such as CITE-sequencing. On the supportive side, a portion of the EPCAM-low or negative, mesenchymal CTCs, were epidermal growth factor receptor (EGFR)-positive, possibly due to the frequent use of anti-EGFR antibodies for CTC positive selection, in combination with anti-EPCAM and anti-ERBB2. Another positive observation, supporting the validity of our work, was that the CTCs from prostate cancer were KLK2 positive, as expected, and conversely all KLK2-positive cells in the integrated datasets were of prostate cancer origin. Thus, notwithstanding the described limitations, we could establish a collection of *bona fide* scRNA-seq CTC profiles from breast cancer after excluding misidentified blood-derived cells and identified novel markers for epithelial and for mesenchymal CTCs, a population which is notoriously difficult to characterize. Such panel of markers will be useful to implement the latest technologies, expanding the spectrum of isolated CTCs, thereby facilitating high-fidelity characterization. Combining multiple targets could improve the capture efficiency, sensitivity, and specificity of CTC isolation. For instance, using both epithelial and mesenchymal markers could help in capturing a broader range of CTCs, including those undergoing EMT and those that may have lost traditional epithelial markers. Another very important result in our work is the design and development of CTCeek a free web-tool for the high-precision CTC annotation (available at https://singlecell.unife.it/ctceek). With this web service, we provide CTC and cancer researchers with a solid instrument to correctly classify scRNA-seq profiles of CTCs. CTCeek was designed to detect and annotate *bona fide* CTCs, separating them from other blood-borne cell types. This goal could be achieved by using our comprehensive reference of PBMC and CTC single-cell profiles. CTCeek is the first computational tool of its kind specifically engineered for the rigorous skimming and classification of CTCs’ scRNA-seq data. Our results confirmed the potential of CTCeek, even in complex settings, i.e. profiles derived from different scRNA-seq technologies. CTCeek demonstrated robust performance that aligned strongly with, and possibly exceeded, two aneuploidy-detection tools, CopyKAT and SCEVAN. Overall, our work advances the understanding of CTC biology and strengthens the foundations for the unbiased and comprehensive investigation of CTCs across different types of solid cancers.

## Methods

Twenty-seven datasets containing RNA profiles from samples annotated as CTCs were obtained from GEO and SRA databases (Supplementary Table 1) and processed to yield a merged expression matrix of 3,302 putative CTC profiles^(17)^. These RNA profiles were generated using two routes, either by scRNA-seq or by classic RNA-seq on single purified cells. A gene was removed from the expression matrix when its counts were not determined for more than 5% of the samples, otherwise undetermined counts were imputed as the median expression for that gene. The single-cell profiles were processed to implement strong quality control (nFeatures >2500 and nCount >10000). The raw counts, obtained after merging the different CTC datasets, were variance stabilized using shifted log^(34)^, the method with the lowest variance after transformation (Supplementary Fig. 9). Heterogeneous CTC samples, likely doublets or multiplets, were identified and removed using scDblFinder^(35)^. The integration of independent datasets was performed using Scanorama^(36)^, and cell cycle genes were removed prior to integration, to avoid influences due to mitotic activity. We identified *bona fide* cancer CTCs in two unrelated ways: i) as cells which were classified as cancer based on their expression profiles, using scATOMIC^(37)^ or ii) as aneuploid cells, using CopyKAT^(27)^. With the goal of performing the most stringent selection towards a “*pure*” single-cell CTC dataset, we also manually curated each putative CTC sample according to its annotation, excluding those passaged in cell culture and those from a multicellular complex. The Broad Institute peripheral blood mononuclear cells (PBMC) Systematic Comparative Analysis (SCA) dataset^(22)^ was used as the reference for blood cells in the integration with CTCs. Various integration methods were assessed, and Scanorama was again chosen as the most balanced method, over FastMNN and Harmony. Comparing the different integration methods, FastMNN was the most aggressive in cluster reduction, but Scanorama led to the lowest number of CTC clusters without over-integration of the PBMC datasets. Tricycle^(38)^ was employed to infer and visualize cell cycle positions. We used the Wilcoxon test coupled with Receiver Operating Characteristic Area Under the Curve (ROC AUC)^(39),(40)^ to select gene markers. R, Bioconductor^(41)^ and RStudio (Posit Software, PBC, Boston, MA, USA) were used for the scRNA-seq analysis described above, alongside Python and Scanpy^(42)^.

The bioinformatic analysis to develop CTCeek was implemented through the development of an interactive web application using the Shiny framework. The primary objective was to provide a user-friendly interface for the automated annotation of CTCs from scRNA-seq data. The underlying algorithm relies on a reference mapping process (anchorage)^(43)^, which identifies cells in a similar biological state between the query and reference datasets (“anchors”) and uses these correspondences to transfer the reference’s cell type labels to each query cell. Crucially, our tool can detect impurities and contaminants (e.g., non-cancer cells) from CTC enrichment experiments, thus overcoming one of the main issues of the common methodologies applied to isolate CTCs^(44)^. Benchmarking statistics, specificity and False Positive Rate, was calculated as:

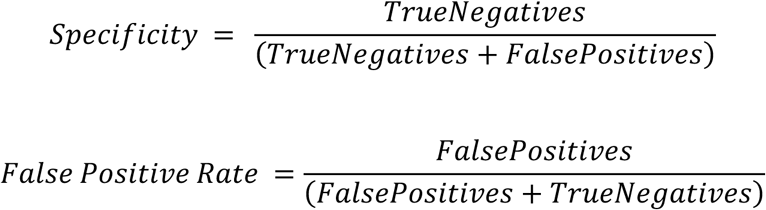

The algorithm detailed methods can be found in the Supplementary Information File, while the scripts are available at https://github.com/PietroAnc/CTCeek.

## Authors’ contributions

Volinia S. conceived and designed the study; Terrazzan A., Ancona P., Carbone F.P., Trevisan P., Szymanek E.A., Palatini J., Kaminski T.S., and Volinia S. collected the data and performed the analysis. Trevisan P., and Ancona P. developed the CTCeek shiny app. Terrazzan A., Ancona P., Carbone F.P., Trevisan P., Zuccato C., Szymanek E.A., Szeląg M., Brugnoli F., Żaczek A., Gaj P., Swiernak M., Calabrò L., Agnoletto C., Palatini J., Bianchi N., Duchnowska R., Senkus-Konefka E., Jazdzewski K., Kaminski T.S., and Volinia S. discussed and revised the methods and results. Terrazzan A., Ancona P., Palatini J., Bianchi N., Senkus-Konefka E., Kaminski T.S., and Volinia S. drafted the manuscript. All the authors read, revised, and approved the final manuscript.

## Data and materials availability

All data needed to evaluate the conclusions in this manuscript are present in the main text and/or in the Supplementary Information File. Processed single-cell gene expression data are available to download at Zenodo (https://doi.org/10.5281/zenodo.17780234).

