## Supplementary File for "Precision single-cell profiling of Circulating Tumour Cells: novel markers and data-driven characterization by CTCeek"

### *SUPPLEMENTARY INFORMATION*

**Supplementary Table 1**. The CTC datasets with the cancer of origin.

| Dataset | N | QC passed | Cancer of Origin |
|---|---|---|---|
| PRJNA471754 | 1100 | 171 | Breast |
| GSE180097 | 293 | 270 | Breast |
| GSE109761 | 289 | 211 | Breast |
| GSE144495 | 195 | 66 | Breast |
| GSE111065 | 172 | 149 | Breast |
| GSE115501 | 136 | 0 | Prostate |
| GSE144494 | 135 | 60 | Breast |
| GSE67980 | 122 | 63 | Prostate |
| GSE117623 | 116 | 0 | Liver |
| GSE144561 | 81 | 0 | Pancreas |
| GSE186288 | 81 | 0 | Breast |
| GSE86978 | 77 | 44 | Breast |
| SRP281893 | 76 | 28 | Melanoma |
| GSE75367 | 74 | 34 | Breast |
| GSE208448 | 68 | 42 | Prostate |
| SRP335264 | 59 | 6 | Colorectal |
| PRJDB11367 | 47 | 20 | Stomach |
| GSE126669 | 31 | 27 | Breast |
| GSE51827 | 29 | 17 | Breast |
| GSE113890 | 27 | 0 | Breast |
| GSE111842 | 16 | 4 | Breast |
| GSE198291 | 16 | 0 | Lung |
| GSE129474 | 15 | 0 | Breast |
| GSE67939 | 15 | 7 | Breast |
| GSE104209 | 12 | 0 | Prostate |
| GSE114704 | 10 | 9 | Pancreas |
| GSE74639 | 10 | 0 | Lung |

**Quality Control (QC)-passed samples are those with a minimal complexity above the QC-thresholds.*

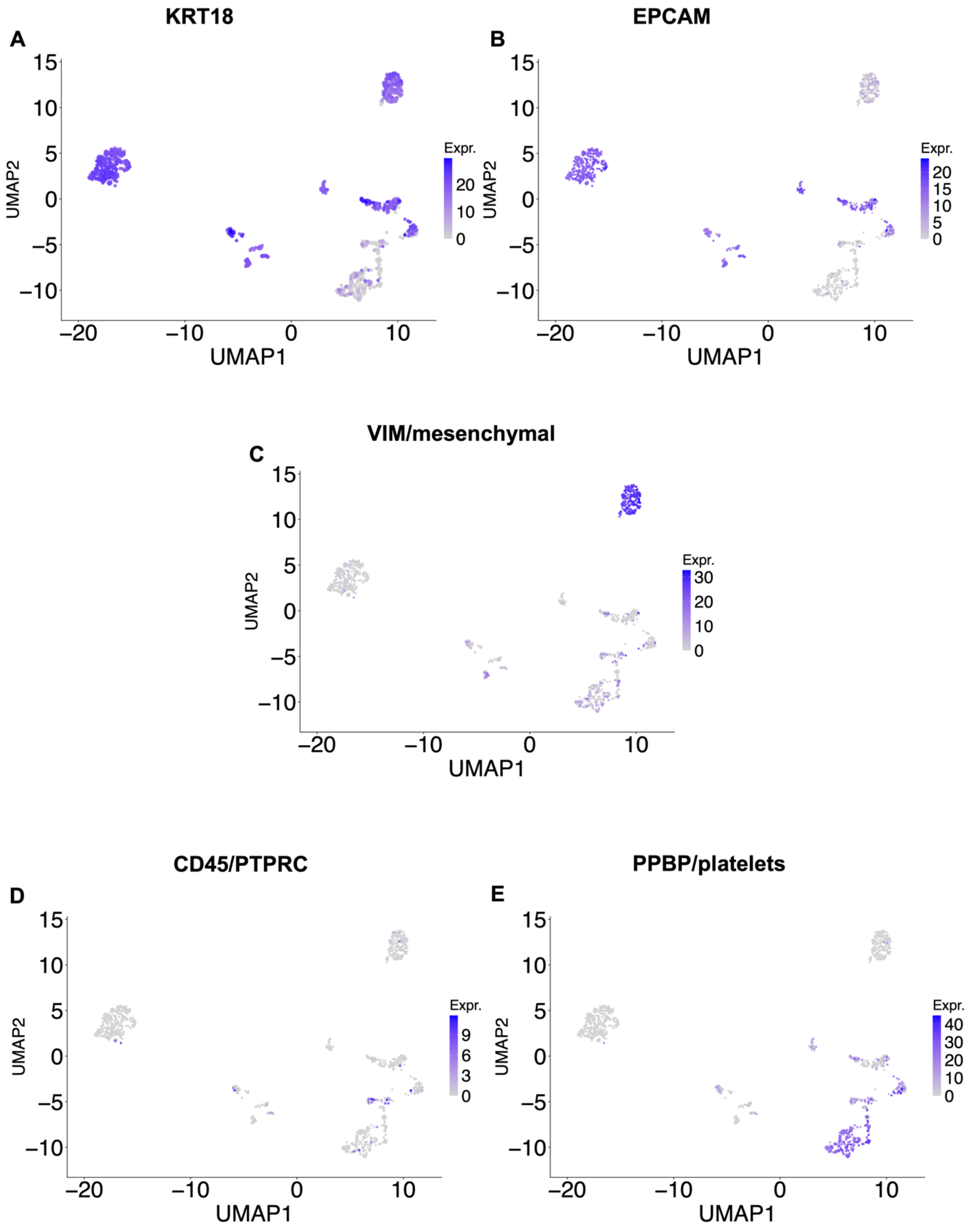


***Supplementary Fig 1***. *Markers' expression plotted over the UMAP of the integrated CTC datasets.*

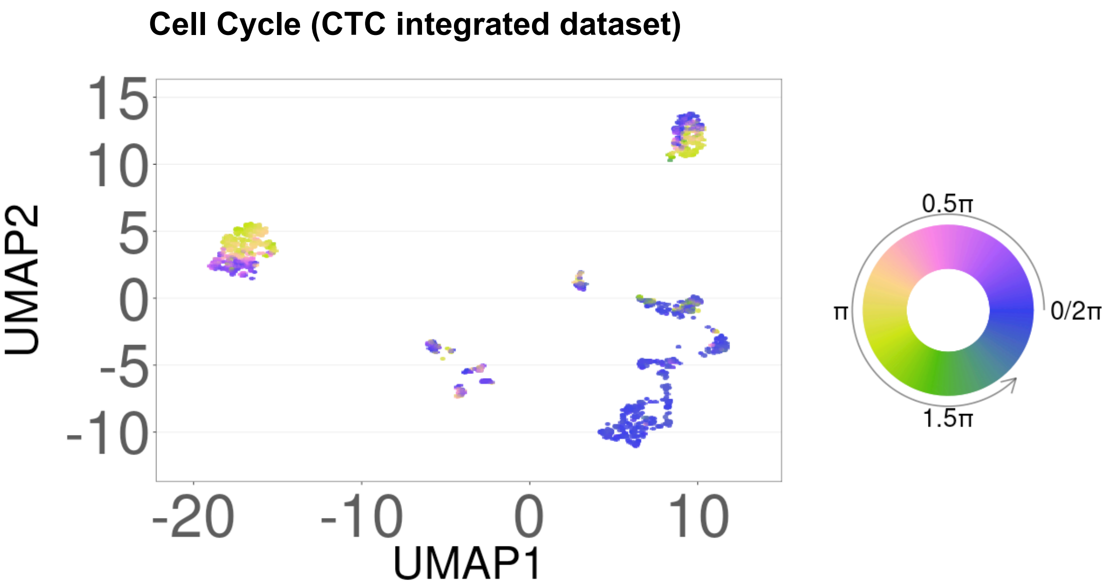


***Supplementary Fig 2.*** *Cell cycle analysis of putative CTCs. Tricycle designs 0.5 pi to be the start of S stage, 0.25 pi to be the start of G2/M stage, 1.5 pi to be the middle of M stage, and 1.75 pi to be G1/G0 stage.*

**Supplementary Table 2.** Differentially expressed genes in CTC contaminant cells.

| Gene | AUC | Avg_log2 FC (sc) | Adjusted p-value | Avg_log2 FC (bulk) | Tissue | Extracellular function |
|---|---|---|---|---|---|---|
| TUBB1 | 0.98 | 21.1 | 0 | 9.8 | platelet | |
| TMEM40 | 0.96 | 21.6 | 0 | 9.4 | platelet | receptor |
| ACRBP | 0.97 | 20.0 | 0 | 9.1 | platelet | |
| SELP | 0.94 | 16.2 | 0 | 9.1 | endothelial | receptor |
| CLEC1B | 0.93 | 14.4 | 0 | 8.7 | platelet | receptor |
| ABCC3 | 0.94 | 26.2 | 0 | 8.6 | | |
| MFAP3L | 0.92 | 21.5 | 0 | 8.6 | | |
| GFI1B | 0.95 | 15.7 | 0 | 8.5 | erythroid | |
| C2orf88 | 0.96 | 21.7 | 0 | 8.5 | | |
| TRIM58 | 0.97 | 22.5 | 0 | 8.5 | erythroid | |
| TAL1 | 0.95 | 15.2 | 0 | 8.3 | | |
| ALOX12 | 0.92 | 17.5 | 0 | 8.3 | platelet | |
| NFE2 | 0.96 | 17.1 | 0 | 8.2 | erythroid | |
| DNM3 | 0.96 | 19.0 | 0 | 8.1 | | |
| PEAR1 | 0.95 | 18.4 | 0 | 8.1 | endothelial | receptor |
| ESAM | 0.94 | 21.6 | 0 | 8.0 | endothelial | receptor |

| | | | | | | |
|---|---|---|---|---|---|---|
| MEIS1 | 0.96 | 13.6 | 0 | 7.7 | | |
| TSPAN33 | 0.91 | 21.3 | 0 | 7.6 | dendritic | receptor |
| GNG11 | 0.95 | 12.4 | 0 | 7.6 | endothelial | |
| PRKAR2B | 0.95 | 13.2 | 0 | 7.5 | | |
| PF4 | 0.94 | 16.1 | 0 | 7.4 | platelet | |
| LGALSL | 0.92 | 11.9 | 0 | 7.2 | | |
| PTGS1 | 0.95 | 13.1 | 0 | 7.2 | | |
| PTCRA | 0.94 | 15.8 | 0 | 7.1 | platelet | |
| GP9 | 0.92 | 17.7 | 0 | 7.1 | platelet | receptor |
| DAB2 | 0.96 | 13.6 | 0 | 7.1 | | |
| SMOX | 0.92 | 15.4 | 0 | 7.1 | platelet | |

**Area Under the Curve (AUC) bulk > 0.99. Benjamini Hochberg adjusted pvalue (BH adj pval) bulk ≤ 0.02. The tissues/markers correspondences were derived from the Panglao DB (https://panglaodb.se/markers.html).*

**Supplementary Table 3.** Contaminant non-cancer cells in the putative CTC datasets.

| Scanorama cluster | Dataset accession | N | CTC purification method |
|---|---|---|---|
| 25 | GSE109761 | 3 | microfluidics and antibody positive selection cocktail |
| 34 | GSE109761 | 1 | microfluidics and antibody positive selection cocktail |
| 25 | GSE111065 | 1 | microfluidics and antibody positive selection cocktail |
| 34 | GSE111842 | 4 | IE/FACS assay with immunomagnetic EpCAM (MJ37) monoclonal antibody (mAb)-coated magnetic beads |
| 25 | GSE144494 | 20 | The CTC-iChip microfluidic device |
| 34 | GSE144494 | 15 | The CTC-iChip microfluidic device |
| 25 | GSE144495 | 10 | The CTC-iChip microfluidic device |
| 34 | GSE144495 | 31 | The CTC-iChip microfluidic device |
| 34 | GSE208448 | 1 | microfluidic-based enrichment of CTCs |
| 25 | GSE51827 | 3 | microfluidics WBC depletion CD45 and CD66b |
| 34 | GSE51827 | 3 | microfluidics WBC depletion CD45 and CD66b |
| 25 | GSE67939 | 2 | microfluidics depletion CD45, CD14 and CD16 |
| 34 | GSE67939 | 3 | microfluidics depletion CD45, CD14 and CD16 |
| 34 | GSE67980 | 4 | CTC-iChip and WBC depletion |
| 25 | GSE75367 | 8 | CTC-iChip and WBC depletion |

| | | | |
|---|---|---|---|
| 34 | GSE75367 | 7 | CTC-iChip and WBC depletion |
| 25 | GSE86978 | 14 | CTC-iChip and WBC depletion |
| 34 | GSE86978 | 10 | CTC-iChip and WBC depletion |
| 25 | PRJDB11367 | 17 | label-free microfluidics |
| 34 | PRJDB11367 | 3 | label-free microfluidics |
| 25 | PRJNA471754 | 134 | label-free microfluidics |
| 34 | PRJNA471754 | 1 | label-free microfluidics |
| 34 | SRP281893 | 28 | label-free microfluidics |
| 25 | SRP335264 | 1 | label-free microfluidics |
| 34 | SRP335264 | 2 | label-free microfluidics |

**The number of non-CTC cells in the contaminant clusters #25 and #34, essentially CD45-negative, KRT-negative, diploid, non-cancer cells which expressed endothelial and platelet markers.*

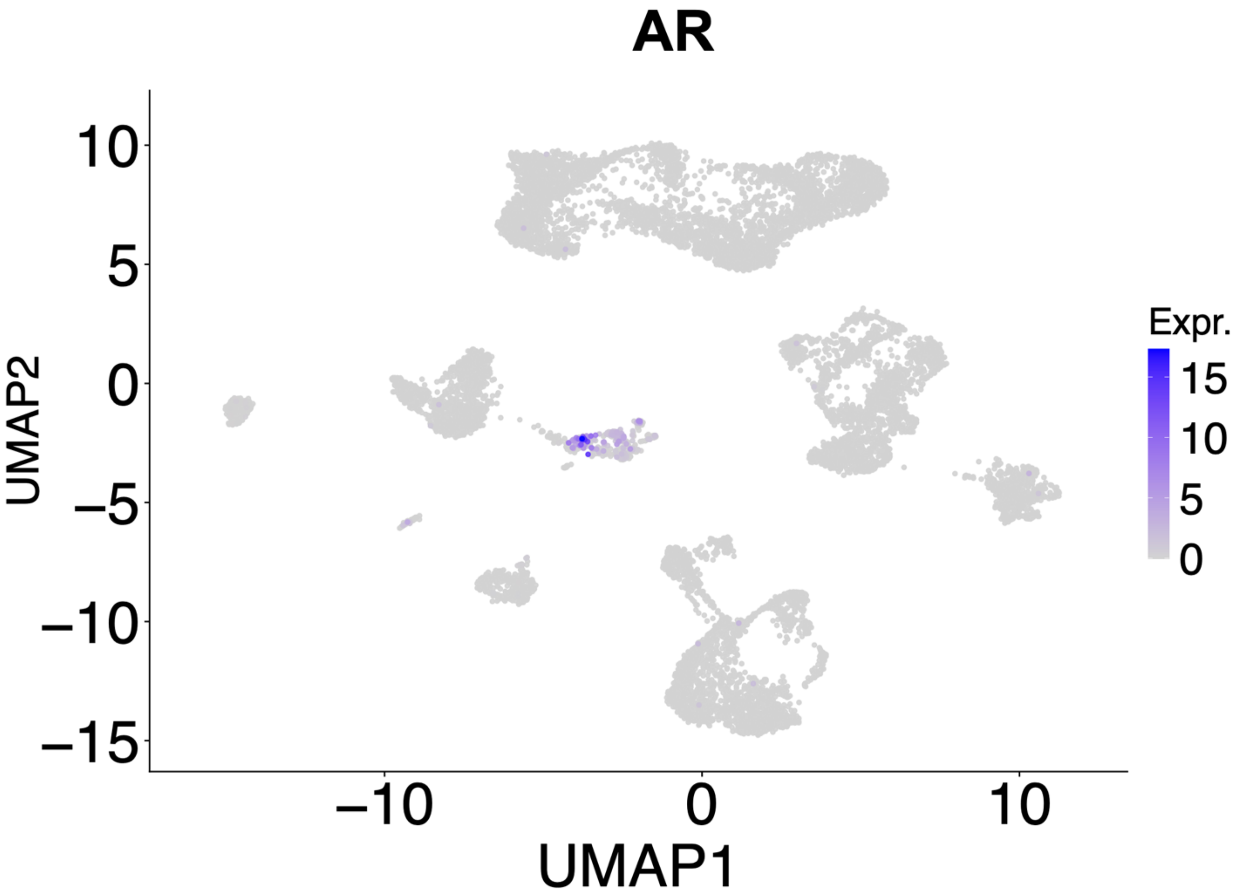


***Supplementary Figure 3.*** *The expression of androgen receptor (AR) is shown over the UMAP plot of integrated CTC and PBMC datasets.*

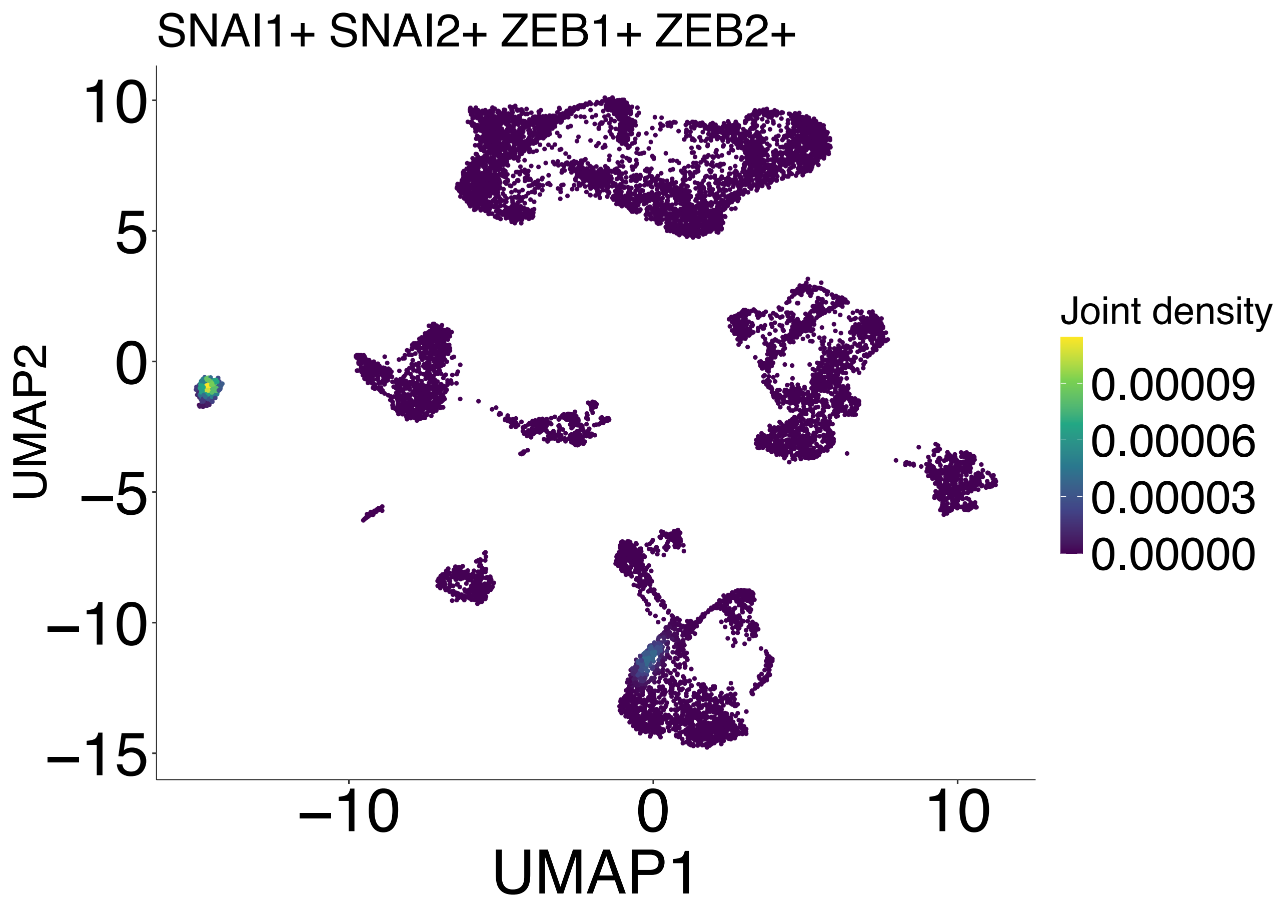


***Supplementary Figure 4.*** *UMAP displaying the cells positive for all four EMT marker genes (SNAI1, SNAI2, ZEB1, and ZEB2. The UMAP plot highlights exclusively the mesenchymal CTC cluster, demonstrating co-expression of EMT genes that define this cell state. Data were analyzed and visualized using the Nebulosa framework to capture joint gene expression density.*

**Supplementary Table 4.** Differentially expressed genes in epithelial A CTCs.

| **Gene** | **AUC (sc)** | **Avg log2 FC (sc)** | **Adjusted p-value (sc)** | **AUC (bulk)** | **Avg log2 FC (bulk)** | **BH adjusted p-value (bulk)** |
|---|---|---|---|---|---|---|
| TACSTD2 | 0.94 | 26.5 | 1.00E-03 | 1.00 | 6.2 | 0.0006 |
| EEF1G | 0.99 | 18.1 | 1.00E-03 | 0.97 | 4.6 | 0.0006 |
| KRT19 | 0.92 | 9.3 | 1.00E-03 | 0.99 | 5.0 | 0.0006 |
| AGR2 | 0.92 | 17.0 | 1.00E-03 | 1.00 | 7.0 | 0.0008 |
| AZGP1 | 0.91 | 18.8 | 1.00E-03 | 1.00 | 8.7 | 0.0028 |

| | | | | | | |
|---|---|---|---|---|---|---|
| CLDN4 | 0.94 | 11.0 | 1.00E-03 | 0.99 | 5.4 | 0.0028 |
| IRF6 | 0.92 | 8.5 | 1.00E-03 | 1.00 | 6.0 | 0.0051 |
| EFNA1 | 0.91 | 10.3 | 1.00E-03 | 0.99 | 4.8 | 0.0053 |
| MLPH | 0.94 | 19.6 | 1.00E-03 | 1.00 | 6.4 | 0.0057 |
| ATP6V0C | 0.92 | 9.2 | 1.00E-03 | 1.00 | 5.2 | 0.0061 |
| DDR1 | 0.93 | 8.7 | 1.00E-03 | 0.99 | 5.1 | 0.0067 |
| HIST1H2BK | 0.93 | 10.6 | 1.00E-03 | 1.00 | 4.9 | 0.0067 |
| MYO6 | 0.92 | 9.5 | 1.00E-03 | 1.00 | 4.1 | 0.0067 |
| HSPB1 | 0.92 | 15.9 | 1.00E-03 | 1.00 | 4.3 | 0.0067 |
| JUP | 0.93 | 11.2 | 1.00E-03 | 0.99 | 4.3 | 0.0067 |
| KRT18 | 0.95 | 8.1 | 1.00E-03 | 0.99 | 4.5 | 0.0067 |
| CD24 | 0.92 | 18.5 | 1.00E-03 | 0.99 | 5.8 | 0.0067 |

**Average log2 Fold Change (Avg log2 FC) bulk >4*

**Supplementary Table 5.** Differentially expressed genes in epithelial B CTCs.

| Gene | AUC (sc) | Avg log2 FC (sc) | Adjusted p-value (sc) | Avg log2 FC (bulk) | BH adjusted p-value (bulk) |
|---|---|---|---|---|---|
| LHX1 | 0.91 | 8.6 | 1.00E-03 | 6.3 | 1.85E-09 |
| ZSCAN1 | 0.93 | 7.6 | 1.00E-03 | 6.1 | 2.07E-08 |
| PKP1 | 1.00 | 10.8 | 1.00E-03 | 8.9 | 4.70E-08 |
| KCNK15 | 0.98 | 8.1 | 1.00E-03 | 7.8 | 1.02E-07 |
| ADCY5 | 0.96 | 7.7 | 1.00E-03 | 7.0 | 1.47E-07 |
| NPNT | 0.99 | 7.5 | 1.00E-03 | 6.3 | 1.47E-07 |
| PPIC | 1.00 | 7.6 | 1.00E-03 | 6.3 | 1.47E-07 |
| GAL | 0.91 | 7.8 | 1.00E-03 | 6.4 | 2.34E-07 |
| PACSIN3 | 0.97 | 7.0 | 1.00E-03 | 6.2 | 2.34E-07 |
| TFAP2C | 1.00 | 7.5 | 1.00E-03 | 6.8 | 2.34E-07 |
| HOXB6 | 0.97 | 7.6 | 1.00E-03 | 6.0 | 4.04E-07 |
| WNT3A | 0.96 | 8.2 | 1.00E-03 | 7.1 | 4.04E-07 |
| HHIPL1 | 0.92 | 9.0 | 1.00E-03 | 7.4 | 7.15E-07 |
| ARHGAP39 | 0.99 | 7.7 | 1.00E-03 | 6.7 | 1.78E-06 |

| | | | | | |
|---|---|---|---|---|---|
| CST6 | 1.00 | 10.7 | 1.00E-03 | 8.1 | 1.78E-06 |
| SP6 | 0.98 | 8.3 | 1.00E-03 | 7.2 | 1.78E-06 |
| LY6K | 0.99 | 7.3 | 1.00E-03 | 6.7 | 9.15E-06 |
| TINAGL1 | 0.99 | 7.8 | 1.00E-03 | 6.6 | 7.44E-05 |
| CRABP2 | 1.00 | 10.6 | 1.00E-03 | 7.4 | 1.69E-04 |
| IGFBP2 | 1.00 | 10.7 | 1.00E-03 | 6.9 | 2.44E-04 |
| ASS1 | 1.00 | 8.2 | 1.00E-03 | 6.2 | 6.98E-04 |
| BMP7 | 0.99 | 7.5 | 1.00E-03 | 7.2 | 1.95E-03 |
| IFI27 | 0.99 | 8.9 | 1.00E-03 | 6.9 | 2.59E-03 |

**Average log2 Fold Change (Avg log2 FC) bulk >6, bulk AUC >0.99*

**Supplementary Table 6.** Differentially expressed genes in all epithelial CTCs.

| Gene | AUC (sc) | Avg log2 FC (sc) | Adjusted p-value (sc) | AUC (bulk) | Avg log2 FC (bulk) | BH adjusted p-value (bulk) |
|---|---|---|---|---|---|---|
| CLDN3 | 0.92 | 20.0 | 1.00E-03 | 1.00 | 7.1 | 2.89E-05 |
| KRT19 | 0.96 | 17.0 | 1.00E-03 | 1.00 | 7.8 | 2.89E-05 |
| S100A14 | 0.91 | 15.0 | 1.00E-03 | 1.00 | 6.7 | 2.89E-05 |
| TACSTD2 | 0.98 | 30.9 | 1.00E-03 | 1.00 | 7.4 | 2.89E-05 |
| SDC1 | 0.96 | 12.9 | 1.00E-03 | 0.99 | 6.7 | 2.89E-05 |
| SPDEF | 0.96 | 13.0 | 1.00E-03 | 1.00 | 6.7 | 4.31E-05 |
| AGR2 | 0.95 | 20.7 | 1.00E-03 | 0.99 | 7.1 | 9.59E-05 |
| EPCAM | 0.99 | 13.2 | 1.00E-03 | 1.00 | 6.6 | 1.00E-03 |
| MAL2 | 0.97 | 12.9 | 1.00E-03 | 1.00 | 6.0 | 1.00E-03 |
| FXYD3 | 0.97 | 13.6 | 1.00E-03 | 0.99 | 6.7 | 1.00E-03 |
| CLDN4 | 0.98 | 20.4 | 1.00E-03 | 1.00 | 7.5 | 1.00E-03 |
| ESRP1 | 0.95 | 12.0 | 1.00E-03 | 1.00 | 6.0 | 1.00E-03 |
| EFNA1 | 0.96 | 18.8 | 1.00E-03 | 1.00 | 7.7 | 1.00E-03 |
| SELENBP1 | 0.96 | 13.4 | 1.00E-03 | 1.00 | 6.7 | 1.00E-03 |
| MLPH | 0.97 | 20.4 | 1.00E-03 | 1.00 | 6.9 | 1.00E-03 |
| CRABP2 | 0.93 | 17.7 | 1.00E-03 | 0.99 | 6.6 | 1.00E-03 |
| MUC1 | 0.93 | 14.2 | 1.00E-03 | 0.99 | 6.7 | 1.00E-03 |

| | | | | | | |
|---|---|---|---|---|---|---|
| KRT18 | 0.99 | 12.4 | 1.00E-03 | 1.00 | 6.3 | 1.00E-03 |
| ELF3 | 0.98 | 12.9 | 1.00E-03 | 1.00 | 6.7 | 1.00E-03 |

**Average log2 Fold Change (Avg log2 FC) single cell (sc) >12, avg_log2 FC bulk >6*

**Supplementary Table 7.** Differentially expressed genes in mitotically active CTCs (epithelial B and mesenchymal).

| **Gene** | **AUC (sc)** | **Avg log2 FC (sc)** | **Adjusted p-value (sc)** | **AUC (bulk)** | **Avg log2 FC (bulk)** | **BH adjusted p-value (bulk)** |
|---|---|---|---|---|---|---|
| TINAGL1 | 0.96 | 9.0 | 1.00E-03 | 1.00 | 6.3 | 0.026 |
| SLCO4A1 | 0.95 | 12.0 | 1.00E-03 | 1.00 | 6.9 | 0.026 |
| ITGB4 | 0.98 | 9.2 | 1.00E-03 | 1.00 | 6.0 | 0.028 |
| LAMA5 | 0.99 | 12.0 | 1.00E-03 | 1.00 | 6.1 | 0.028 |
| TUBA1C | 0.97 | 9.4 | 1.00E-03 | 1.00 | 4.3 | 0.028 |
| IGFBP4 | 0.93 | 15.1 | 1.00E-03 | 0.99 | 6.2 | 0.028 |
| CRABP2 | 0.90 | 9.9 | 1.00E-03 | 0.96 | 5.9 | 0.028 |
| PYGB | 0.96 | 10.5 | 1.00E-03 | 0.98 | 4.5 | 0.028 |
| SPINT2 | 0.95 | 10.9 | 1.00E-03 | 0.95 | 4.0 | 0.035 |

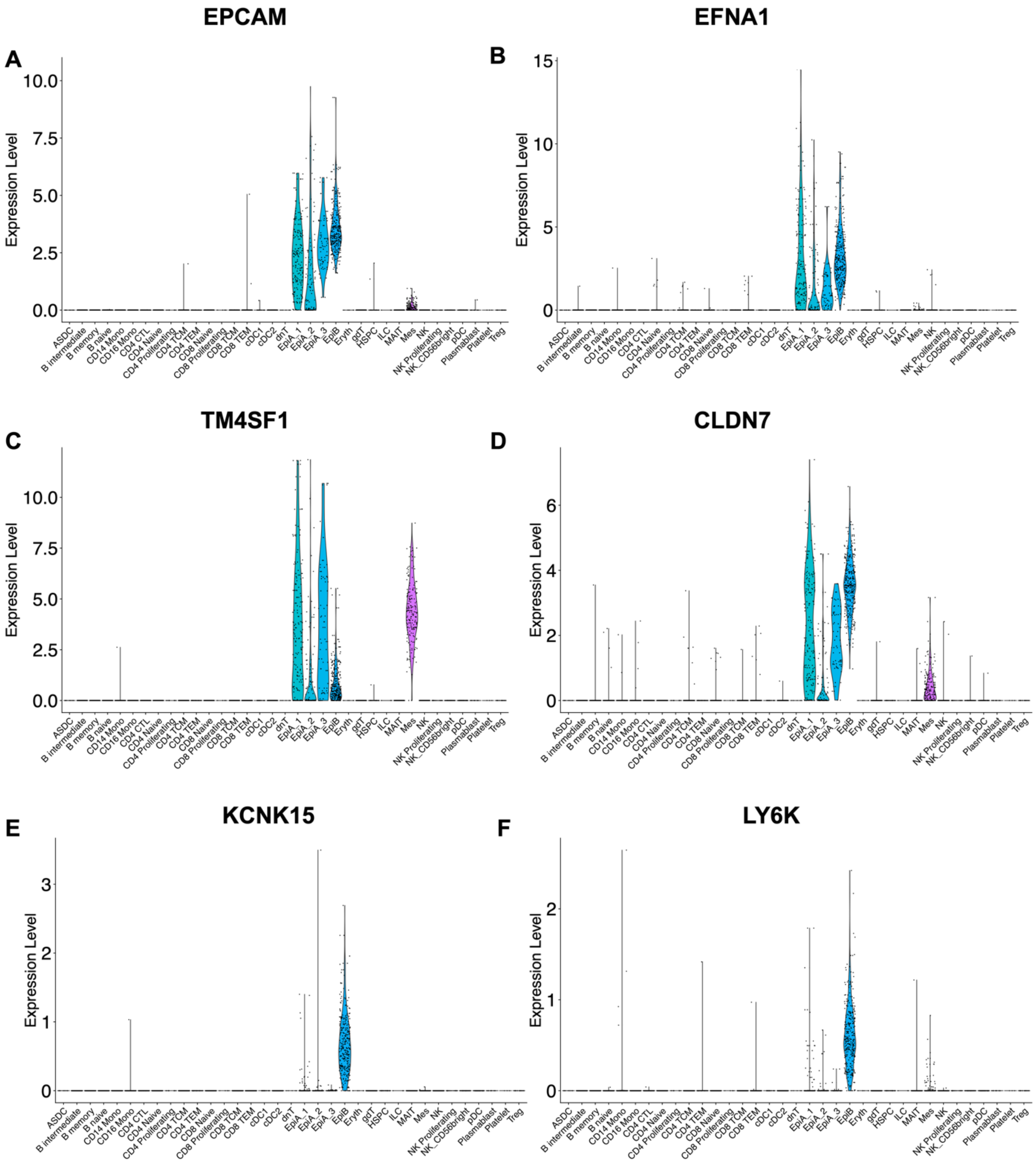


***Supplementary Figure 5.*** *The expression of various CTC markers shown for the cell types in the integrated CTC and PBMC datasets. (**A**) EPCAM, the current CTC gold standard; (**B**) Expression of EFNA1, a novel marker for epithelial CTCs; (**C** and **D**) TM4SF1 and CLDN7 expressions, which are the new markers for both epithelial and mesenchymal CTCs. Finally, the expression of KCNK15 and LY6K, two new specific markers for epithelial B CTCs, are highlighted in panels **E** and **F** respectively.*

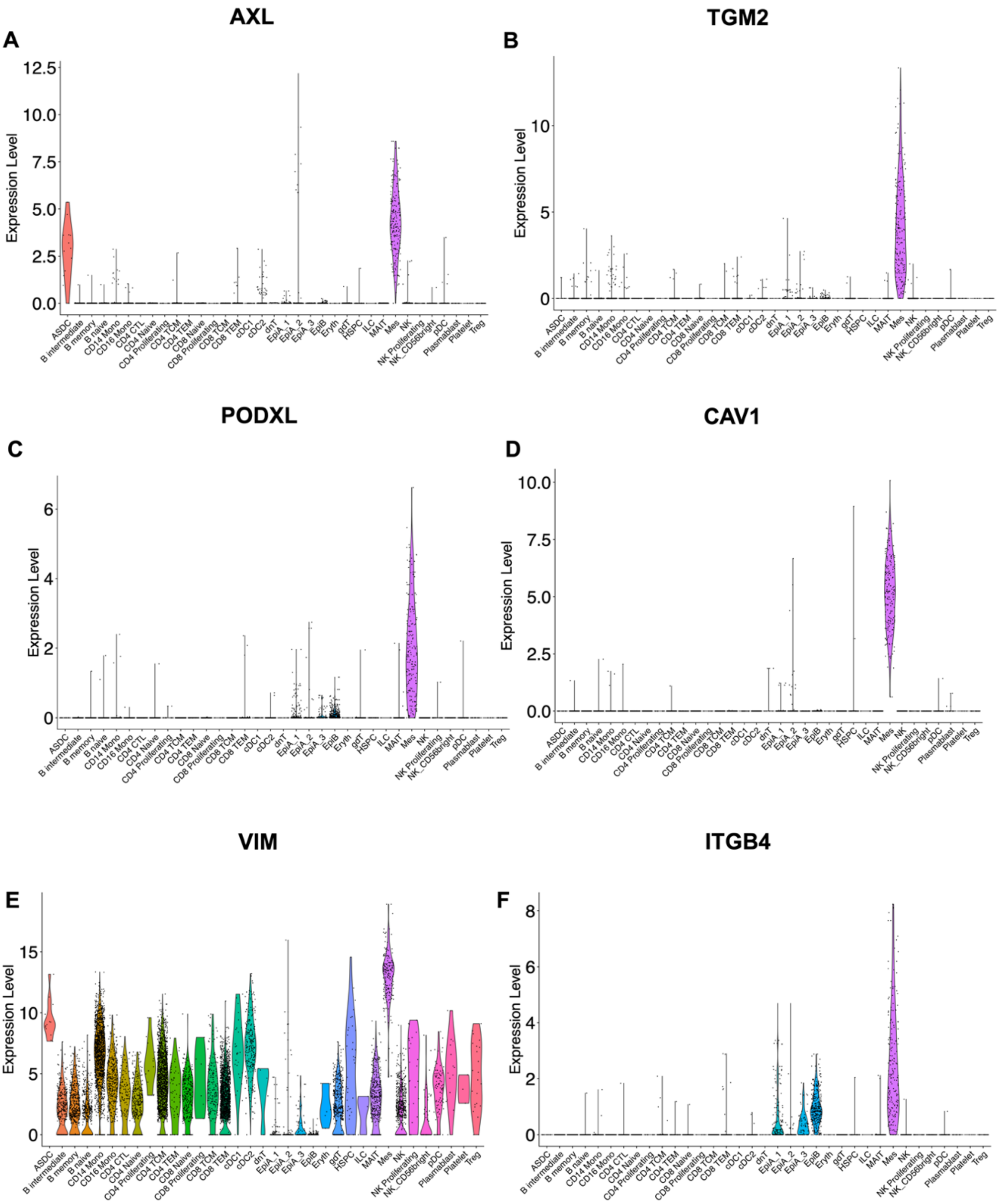


***Supplementary Figure 6.*** *Different expression of the newly mesenchymal markers, shown for the cell types in the integrated CTC/PBMC datasets. (**A**) AXL is a marker for mesenchymal CTCs and ASDCs; the expression of TGM2, PODXL, and CAV1 novel markers for mesenchymal CTCs is depicted in panel **B**, **C** and **D**; (**E**) Expression of VIM, a classical marker for mesenchymal cells, but not specific for CTCs; (**F**) ITGB4, a component of the integrin receptor that binds to laminin, is overexpressed in epithelial B and mesenchymal CTCs, which are the CTCs with the highest engagement in cell cycle.*

**Supplementary Table 8.** Differentially expressed genes in PBMC compared with CTCs.

| **Gene** | **AUC (sc)** | **Avg log2 FC (sc)** | **Adjusted p-value (sc)** | **AUC (bulk)** | **Avg log2 FC (bulk)** | **BH adjusted p-value (bulk)** |
|---|---|---|---|---|---|---|
| CD52 | 0.95 | 17.7 | 0.00E+00 | 1.00 | 6.1 | 4.79E-04 |
| PTPRC | 0.91 | 13.8 | 0.00E+00 | 1.00 | 5.5 | 4.79E-04 |
| CXCR4 | 0.81 | 12.7 | 1.37E-220 | 1.00 | 5.5 | 4.79E-04 |
| CD37 | 0.88 | 13.0 | 0.00E+00 | 1.00 | 5.2 | 4.79E-04 |
| CD48 | 0.85 | 12.6 | 1.46E-278 | 1.00 | 4.8 | 4.79E-04 |
| HCST | 0.81 | 13.0 | 2.31E-215 | 1.00 | 4.5 | 4.79E-04 |

**Average log2 Fold Change (Avg log2 FC) single cell (sc) >12, avg_log2 FC bulk >4*

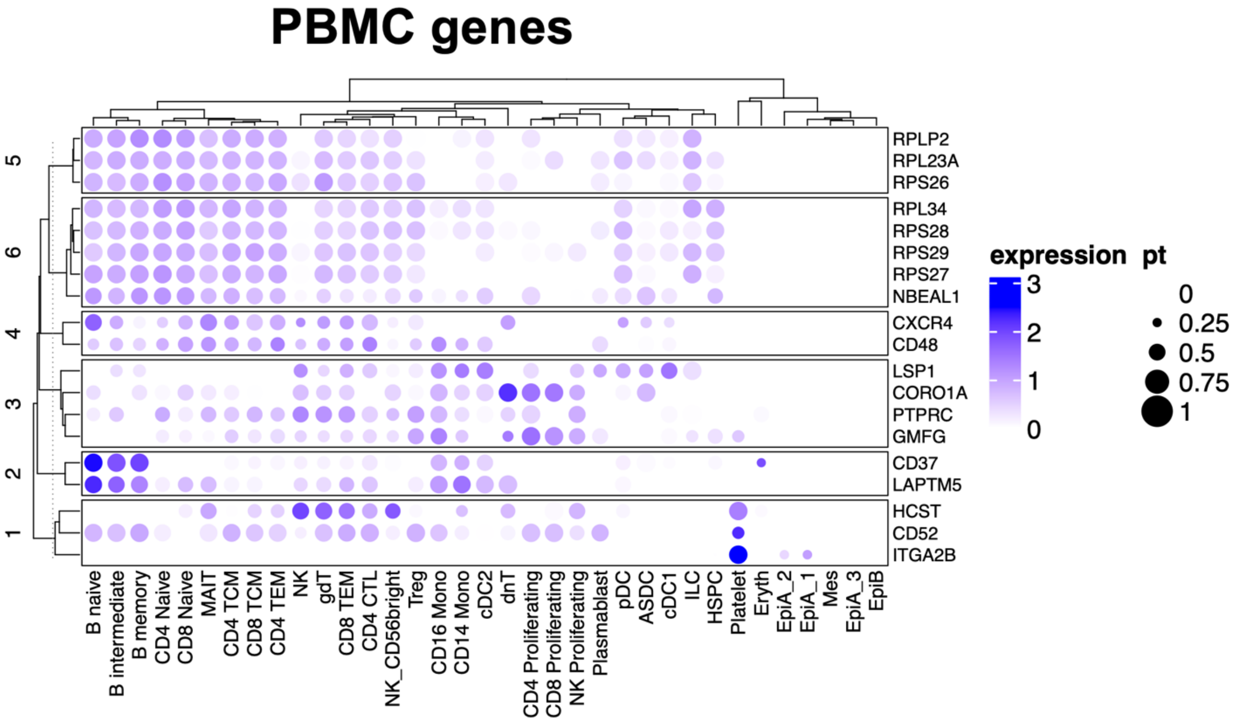


***Supplementary Figure 7.*** *Markers for negative selection of hematopoietic cells. Clustered dotplot with the genes over-expressed in PBMC cells after microfluidics-based size selection and CTC enrichment. CTC subgroups are indicated together with the hematopoietic predicted cell types. The radius of the circle is proportional to the percent (pt) of positive cells in each lane.*

**Supplementary Table 9.** Differentially expressed genes in ASDCs compared with CTCs.

| Gene | AUC | Avg log2 FC | Adjusted p-value |
|---|---|---|---|
| TYROBP | 1.00 | 11.5 | 7.07E-41 |
| HLA-DRB1 | 0.99 | 14.4 | 7.15E-14 |
| IRF8 | 0.97 | 11.8 | 4.07E-52 |
| LSP1 | 0.89 | 10.8 | 3.59E-16 |
| CD48 | 0.87 | 10.7 | 2.49E-38 |
| WAS | 0.87 | 9.8 | 2.25E-21 |
| PTPRC | 0.86 | 9.6 | 8.91E-12 |

**Supplementary Table 10.** Patients' information from the validation datasets.

| Cancer type | N of patients | N of CTCs | Cohort |
|---|---|---|---|
| Prostate cancer | 3 | 108 | GSE255889; GSE295441 |
| Pancreatobiliary cancer | 2 | 7 | GSE295441 |
| Non-small cell lung cancer | 1 | 6 | GSE295441 |
| Breast cancer | 1 | 10 | GSE295441 |
| Colorectal cancer | 2 | 5 | GSE295441 |

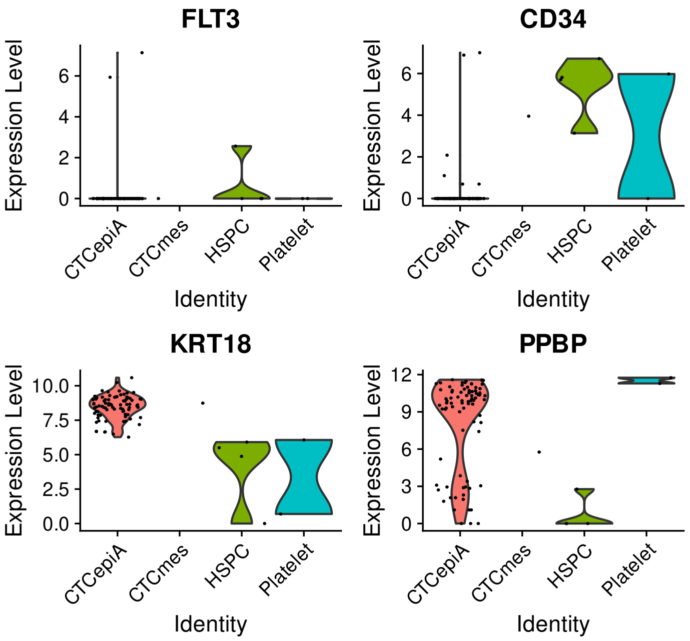


***Supplementary Figure 8****. Validation of CTCeek predictions through marker-based discrimination of cell identities. Violin plots of lineage-defining marker expression (FLT3, CD34, VWF, KRT18, PPBP) across predicted populations. Marker distributions validate CTCeek's single-cell classification and resolve discordant predictions through biological marker concordance.*

**Supplementary Table 11.** CTCeek performance comparison against CNV state-of-art tools.

| Comparison | N | Agreement | Concordance rate | CTCeek CTCs | Reference CTCs | Shared CTCs | Jaccard index[a] |
|---|---|---|---|---|---|---|---|
| CTCeek *vs* CopyKAT | 136 | 117 | 86.03% | 93 | 110 | 92 | 0.8288 (82.88%) |
| CTCeek *vs* SCEVAN | 136 | 119 | 87.5% | 93 | 88 | 82 | 0.8283 (82.83%) |
| CTCeek *vs* double-positives | 136 | 82 | 93.18% | 93 | 88 | 82 | 0.9318 (93.18%) |

[a] *The Jaccard Index has been calculated using the formula:*

$$J(A,B) = \frac{|Inter\sec t\,ion|}{|Union|} = \frac{Shared\ positive\ cells\ (CTCs)}{Total\ unique\ positive\ cells\ (CTCs)}$$

### *SUPPLEMENTARY METHODS*

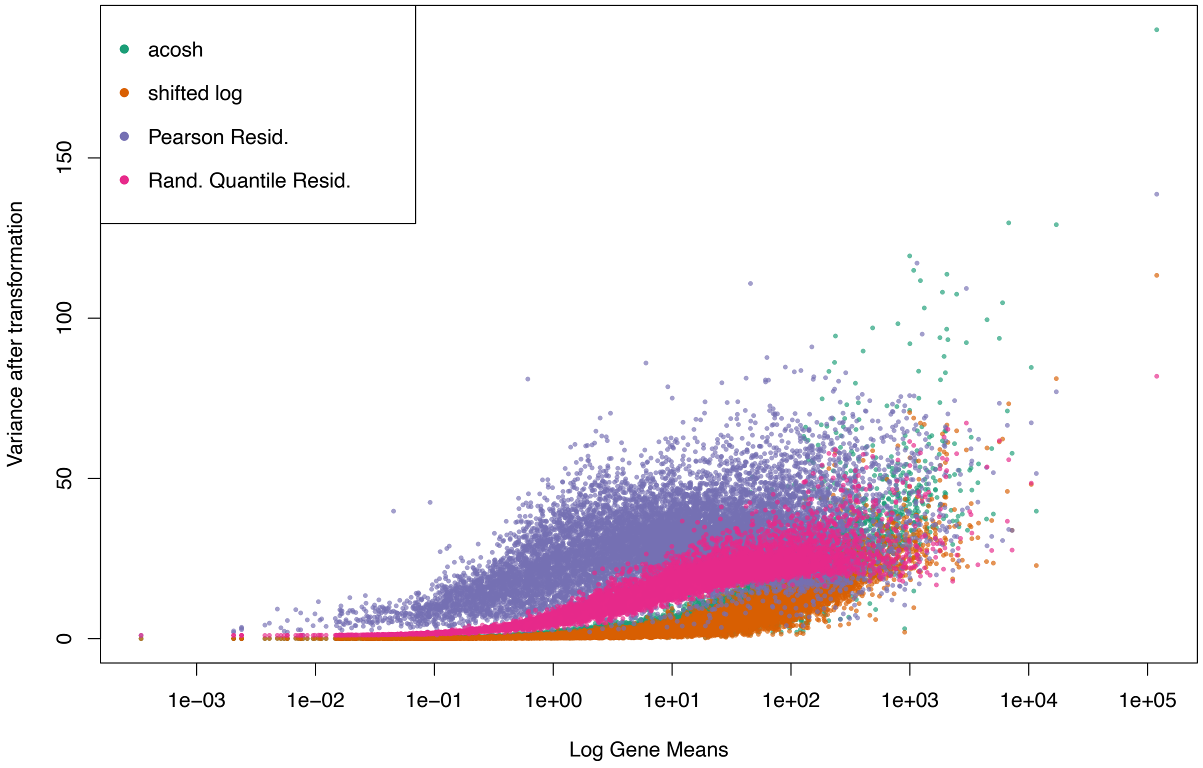


***Supplementary Fig 9.*** *Performance of different data transformation methods.*

#### CTCeek pipeline for algorithm building

The bioinformatic analysis to develop CTCeek was implemented through the development of an interactive web application using the Shiny framework. The primary objective was to provide a user-friendly interface for the automated annotation of CTCs from scRNA-seq data. The application relies on a suite of essential R packages, including shiny, shinyjs, shinycssloaders, shiny.emptystate, Seurat, dplyr, DT, ggplot2, patchwork, future, and promises, ensuring robust performance and responsiveness for large datasets.

The application requires a Seurat object, saved in *.rds* format, as input, which is directly uploaded by the user via the interface. As first step, the tool automatically performs quality control check over the user-uploaded file, plotting the nCounts, nFeatures and the percentage of mitochondrial genes.

Then, if the SCTransform assay is not present, the expression data are subjected to normalization and variance stabilization using the SCTransform algorithm, implemented via the SCTransform function in the Seurat package. Then dimensionality reduction is performed using Principal Component Analysis (PCA). The number of principal components ($_N$PCs) utilized is determined dynamically by an internal function (safe_npcs), which ensures an optimal number is selected, avoiding the use of more components than supported by the dataset's dimensions or a predefined maximum (e.g., $_N$PCs≤50). Cellular annotation is performed using a label transfer approach based on a pre-processed, locally stored Seurat reference dataset. This reference object contains previously validated cell type labels and a pre-computed UMAP embedding. Correspondences between the experimental dataset (query) and the reference dataset are identified using the FindTransferAnchors function in Seurat, which calculates inter-dataset relationships in the reduced dimensional space. The MapQuery function is subsequently applied to project the query cells onto the reference UMAP embedding, facilitating co-visualization and label transfer. Each query cell is assigned a predictive cell identity (highest prediction score) based on the reference. The predicted cell labels are stored within the metadata of the query Seurat object under the field "CTCeek_final_prediction".

The application provides a summary count of identified CTCs across the classes of interest, namely CTCepiA, CTCepiB, and CTCmes, while annotating the contaminant cells to the respective blood-derived cell type. An auxiliary analytical module allows for the identification of cluster-specific marker genes using a one-*vs*-all differential expression approach. Differentially Expressed Genes (DEGs) are calculated using the FindMarkers function in Seurat (using non-parametric Wilcoxon rank sum test), comparing gene expression in a user-selected cell group (cluster or annotation) against all remaining cells combined. The analysis yields genes that are significantly upregulated in the target cluster, typically filtered based on log-fold change and adjusted *p*-value. The graphical interface was specifically customized to enable side-by-side visualization of the reference dataset's UMAP and the projected experimental cells (Supplementary Fig. 10). All generated plots are exportable in high-resolution *.png* format. Intermediate and final processing steps (pre-processing, UMAP projection, marker calculation) are shared to the user via an integrated notification system. Analysis results, including cell annotations and marker gene lists, are presented in interactive tables, and are available for download in *.csv* format. The final annotated Seurat object can also be exported in *.rds* format for downstream analysis. All the necessary files to reproduce our benchmarking and run CTCeek locally can be found on the corresponding github directory (https://github.com/PietroAnc/CTCeek).

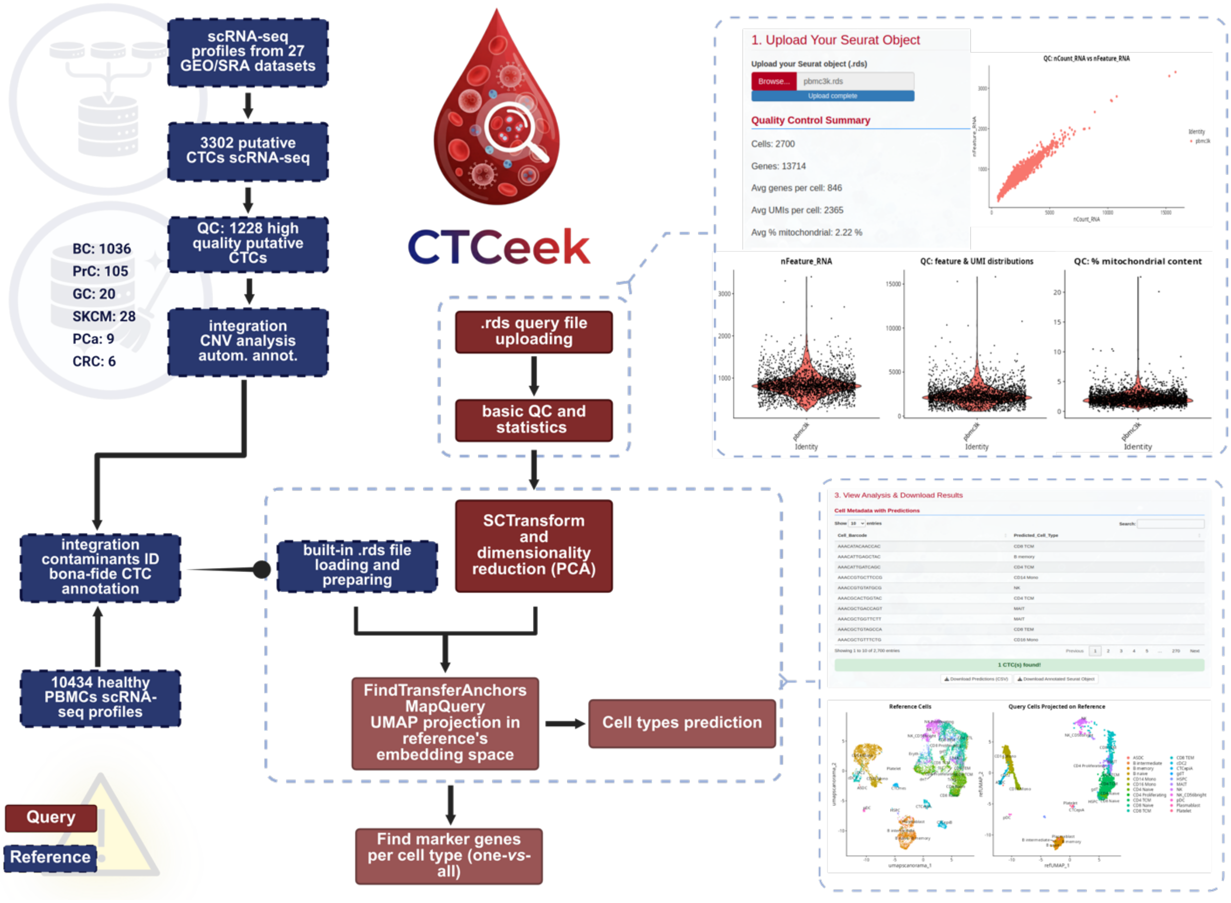


***Supplementary Fig 10.*** *CTCeek framework. The workflow demonstrates the complete analysis process from .rds object upload through quality control, data transformation (SCTransform and PCA), cell type prediction using transfer learning (FindTransferAnchors and MapQuery), and identification of marker genes. Results include QC plots, predicted cell types with marker genes, UMAP projections showing reference and query cell populations, and dimensional reduction visualizations.*
